# IL-21 signaling promotes the establishment of KSHV infection in human tonsil lymphocytes by increasing early targeting of plasma cells

**DOI:** 10.1101/2021.12.06.471359

**Authors:** Nedaa Alomari, Farizeh Aalam, Romina Nabiee, Jesus Ramirez Castano, Jennifer Totonchy

## Abstract

Factors influencing Kaposi’s sarcoma-associated herpesvirus (KSHV) transmission and the early stages of KSHV infection in the human immune system remain poorly characterized. KSHV is known to extensively manipulate the host immune system and the cytokine milieu, and cytokines are known to influence the progression of KSHV-associated diseases. Here, using our unique model of KSHV infection in tonsil lymphocytes, we investigate the influence of host cytokines on the establishment of KSHV infection in human B cells. Our data demonstrate that KSHV manipulates the host cytokine microenvironment during early infection and susceptibility is generally associated with downregulation of multiple cytokines. However, we show that IL-21 signaling promotes KSHV infection by promoting both plasma cell numbers and increasing KSHV infection in plasma cells as early as 3 days post-infection. Our data reveal that this phenotype is dependent upon a specific milieu of T cells, that includes IL-21 producing Th17, Tc17 and CD8+ central memory T cells. These results suggest that IL-21 plays a significant role in the early stages of KSHV infection in the human immune system and that specific immunological states favor the initial establishment of KSHV infection by increasing infection in plasma cells.

## Introduction

Kaposi’s Sarcoma Herpesvirus (KSHV) is a lymphotropic gamma-herpesvirus, originally discovered as the causative agent of Kaposi Sarcoma (KS) [1]. KS is a highly proliferative tumor derived from lymphatic endothelial cells [2]. KSHV is also associated with the B cell lymphoproliferative diseases, Primary Effusion Lymphoma (PEL) and Multicentric Castleman’s Disease (MCD) [3, 4], as well as the inflammatory disorder KSHV inflammatory cytokine syndrome (KICS) [5]. KSHV is linked to 1% of all human tumors, and the World Health Organization (WHO) has classified it as class I carcinogen [6, 7]. KSHV infection is asymptomatic in most healthy individuals, and KSHV-associated malignancies arise primarily in immunocompromised patients. Indeed, KS remains one of the most common cancers in people living with HIV/AIDS [8].

The geographical distribution of KSHV is not ubiquitous. KSHV infection is endemic in sub-Saharan Africa and in the Mediterranean basin. KSHV prevalence is also high in subpopulations in other parts of the world such as men who have sex with men (MSM). Saliva is the only secretion where KSHV DNA is commonly detected [9], and, based on this, person-to-person transmission of KSHV is thought to occur via saliva. The oral lymphoid tissues are rich in KSHV target cell types including lymphatic endothelial cells and B cells, and are therefore a likely site for the initial establishment of KSHV infection in a new human host. However, the exact mechanisms for KSHV transmission and how environmental, behavioral and host factors influence transmission and early infection events remain to be definitively established. This gap in our understanding dramatically affects our ability to find efficient strategies to decrease the transmission or influence host-level susceptibility to KSHV infection. We previously analyzed susceptibility to KSHV infection in a cohort of human tonsil samples with diverse race, sex and age distributions and found that these samples displayed high variability in susceptibility that could not be linked to demographic factors [10]. Our ongoing research seeks to identify, and mechanistically characterize, host-level susceptibility factors that influence this variable susceptibility. It is important to note that, in this context, it is the highly susceptible and highly refractory “outlier” specimens that may ultimately prove the most informative in identifying these critical susceptibility factors.

Cytokine dysregulation is strongly linked to the pathogenesis of KSHV-associated lymphoproliferations [11]. However, their contribution to the early stages of KSHV infection and whether the cytokine milieu in the oral cavity contributes to host-level susceptibility to KSHV infection is unclear. IL-21 is a pleiotropic cytokine that has diverse effects on B cell, T cell, macrophage, monocyte, and dendritic cell biology. It is produced mainly by natural killer T (NKT) cells and CD4+ T cells, including follicular helper (Tfh) cells [12]. The IL-21 receptor is expressed by several immune cells, including B and T cells and is comprised of a unique IL-21R subunit and the common cytokine receptor *γ* chain (CD132), which is also part of the receptor for IL-2, IL-4, IL-7, IL-9, and IL-15 [13]. IL-21 plays a critical role in B cell activation and expansion [14], as well as B cell differentiation to immunoglobulin (Ig)-secreting plasma cells. The regulation of maturation of B cells into plasma cell is driven by the several transcription factors including Blimp1 and Bcl6 [15], which can both be induced by IL-21 signaling, indicating that IL-21 is an important regulator of plasma cell differentiation [16, 17]. IL-21 has been studied in the pathogenesis of chronic lymphocytic choriomeningitis virus (LCMV) infection, influenza virus, and, most relevant to this study, IL-21 plays an important role in the early establishment of murine gammaherpesvirus 68 (MHV68) infection in mice [18-20]. Moreover, IL-21 induces differentiation of B-lymphoblastoid cell lines into late plasmablast/early plasma cell phenotypes, and regulates the expression of many latent proteins in EBV^+^ Burkitt lymphoma cell lines [21, 22]. Although IL-21 is canonically thought to act primarily in germinal centers, it is detected in interfollicular areas in MCD patients [23]. There are few studies, to date, examining the contribution of IL-21 to KSHV infection and KSHV-associated disease.

In this study, we use our well-established *ex vivo* tonsil lymphocyte infection model to explore whether KSHV alters cytokine secretion early in infection and whether cytokine levels have an effect on the establishment of KSHV infection. As with our previous studies exploring factors influencing susceptibility of tonsil samples to KSHV infection, we concentrated data collection on 3 dpi, as the earliest timepoint in which infection can be reliably detected, in order to maximize our ability to identify intrinsic susceptibility factors and minimize the contribution of cell culture artifacts that accumulate over time in the cultures. We identify IL-21 as a factor that specifically influences KSHV infection in plasma cells, which we previously characterized as a highly targeted cell type in early infection [10]. We show that supplementation of tonsil lymphocyte cultures with IL-21 enhances total infection while IL-21 neutralization decreases total KSHV infection at early times post-infection. We demonstrate that IL-21 signaling and KSHV infection synergistically increase plasma cell frequencies in the cultures, there are increased levels of KSHV-infected plasma cells in the presence of IL-21, and that these effects are correlated with total KSHV infection in all B cell subsets at 3 dpi. We further explore the immunological mechanisms of this IL-21 effect by establishing which B cell types are responding IL-21, and what T cell subsets are producing IL-21 in our model.

These results identify IL-21 signaling as a factor that influences the establishment of KSHV infection in B lymphocytes via manipulation of plasma cells. Together with our previous work, this study underscores the importance of plasma cell biology in the initial establishment of KSHV infection in the oral lymphoid tissues. Moreover, we identify a combination of T cells, including particular IL-21 secreting T cell subsets, that correlate with KSHV-mediated manipulation of plasma cells. Thus this T cell signature may be indicative of an inflammatory state that favors KSHV transmission.

## Results

### Host cytokines influence the establishment of KSHV infection B lymphocytes

Despite the critical interplay between KSHV and host cytokine signaling, little is known about whether cytokines influence host susceptibility to KSHV infection. In fact, the roles of proinflammatory cytokines during KSHV infection have been studied mostly in naturally-infected human B cell lines derived from PEL [24, 25]. In order to examine whether cytokines alter the early stages of KSHV infection in the tonsil, we quantitated the levels of 13 cytokines in the supernatants of Mock and KSHV-infected tonsil lymphocyte cultures at 3 days post-infection using a bead-based multiplex immunoassay. This dataset includes 33 independent infections using 24 unique tonsil specimens. IL-6, IFNγ, TNFα and IL-22 were the most prevalent cytokines in our cultures based on the median values overall (Fig 1A, panel order). IL-6 was the only cytokine significantly induced in KSHV-infected cultures compared to Mock cultures (Fig 1A) and the magnitude of IL-6 induction by KSHV infection is far greater than the effect of infection on any other cytokine (Fig 1B). This result is consistent with our previous results in cultures containing only naïve B lymphocytes [26]. Our data also reveals statistically significant reductions in IL-5 and IL-4 concentrations in KSHV-infected cultures compared to Mock cultures (Fig 1A), but these changes are very small compared to those seen with IL-6 (Fig 1B). Interestingly, IFNg concentrations were highly affected by KSHV infection, but there were sample-specific differences in whether this effect was positive or negative (Fig 1B).

**Figure 1:**
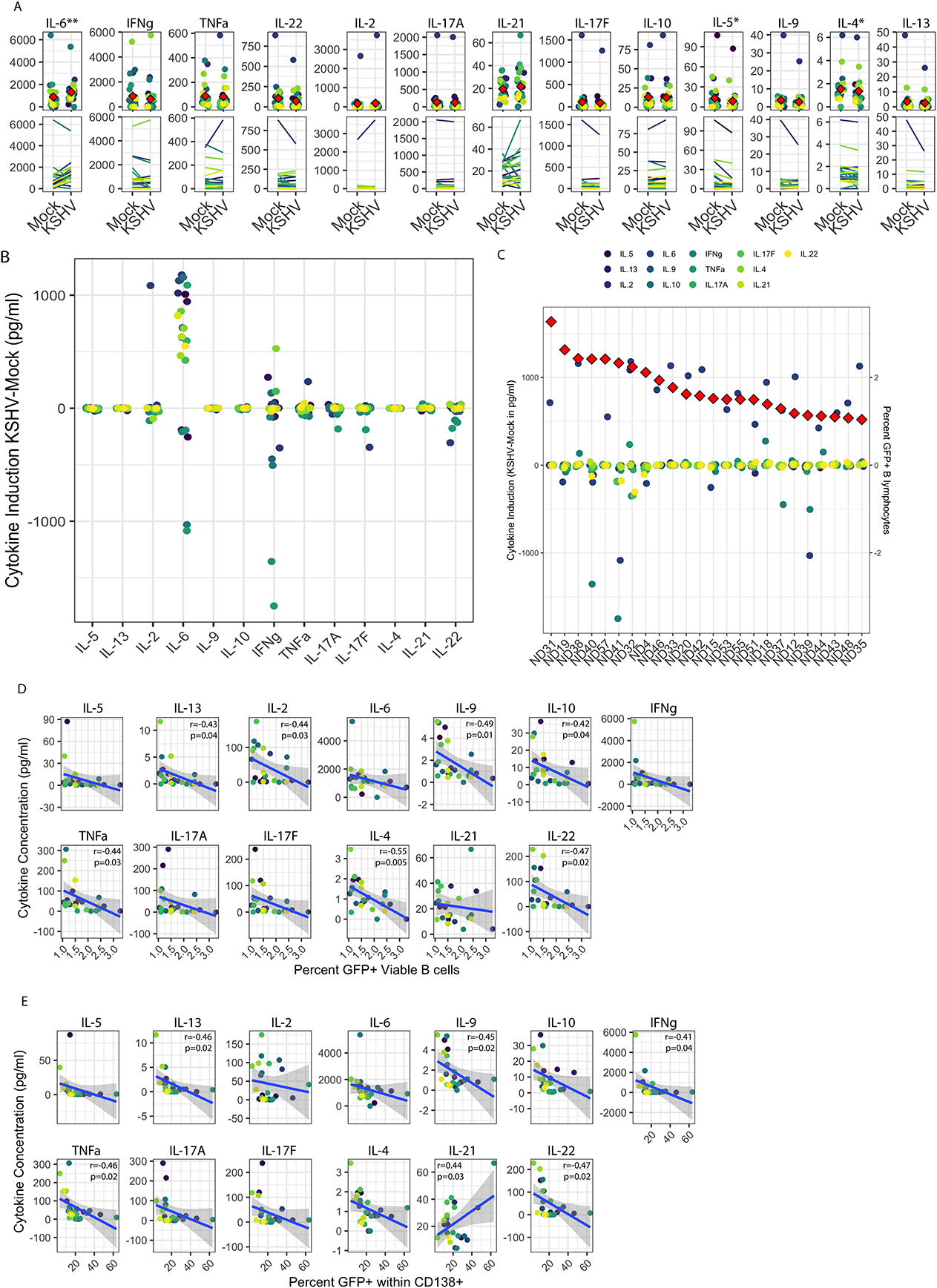
KSHV alters cytokine secretion and cytokines affect the establishment of infection. 33 replicate infections using 24 unique tonsil specimens were performed using KSHV.219 infection of naïve B lymphocytes followed by reconstitution of the total lymphocyte environment and culture on CDW32 feeder cells. At 3 dpi, cells were collected for flow cytometry analysis for infection (GFP) and B cell subsets using our previously-characterized immunophenotyping panel and supernatants were collected for analysis of cytokines by multiplex immunoassay (Biolegend Legendplex) **(A)** cytokine production in Mock and KSHV-infected cultures showing individual sample quantities and means (red diamonds, top panels) and matched Mock and KSHV samples to show trends of induction/repression (bottom panels) Statistical analysis was performed by one-way repeated measures ANOVA. p=0.01 F=7.06 for IL-5, p=0.0001 F=14 for IL-6, p=0.5 F=4.3 for IL-4 **(B)** Data as in (A) showing the level of induction or repression of each cytokine comparing KSHV to matched Mock cultures **(C)** Induction or repression of all cytokines (left y-axis) on a per-sample basis ordered based on overall susceptibility based on percentage of GFP+ B lymphocytes in the same culture (right y-axis, red diamonds) **(D)** Pairwise correlations using Pearson method between cytokine concentration (y-axis) and overall infection (x-axis) in KSHV-infected lymphocyte cultures **(E)** Pairwise correlations using Pearson method between cytokine concentration (y-axis) and Percent GFP+ within CD138+ (x-axis) in KSHV-infected lymphocyte cultures. For panels A, B, D and E colors indicate individual tonsil specimens and can be compared between panels. For D and E, significant correlations are indicated with r and p-values on the individual panels.

In order to determine whether cytokines affect the establishment of KSHV infection, we examined whether the concentration of cytokines in the supernatants of KSHV-infected cultures is correlated with the level of infection in B lymphocytes (based on GFP reporter expression) in the same culture by flow cytometry analysis (Fig 1C & D). On a per-sample level, many individual cytokines (notably IL-6 and IFNg) were induced or repressed independent of susceptibility (Fig 1C, red diamonds and right y-axis scale). However, many of the most susceptible samples included in this dataset, display lower levels of multiple cytokines in KSHV-infected cultures vs. Mock cultures (Fig 1C, far left samples), indicating that the ability of KSHV to suppress cytokine secretion may influence the early stages of infection. Consistent with this, pairwise comparisons between overall GFP level in B lymphocytes and the level of each cytokine in the KSHV-infected cultures revealed universally negative correlations between overall KSHV infection and cytokine levels with lower cytokine levels observed in more susceptible samples. These negative correlations were statistically significant for IL-2, IL-9, IL-10, TNFα, IL-4 and IL-22 (Fig 1D). Since plasma cells were identified as a highly targeted cell type in our previous study [10], we examined the correlations between cytokine levels in KSHV-infected cultures and infection in the CD138+ plasma cell subset. This analysis revealed negative correlations similar to those seen with overall infection with IL-13, IL-9, IFNγ, TNFα and IL-22 levels showing statistically significant negative correlations with plasma cell infection. However, in this analysis IL-21 levels showed a significant positive correlation with plasma cell infection (Fig 1E). Taken together, these data demonstrate that (1) KSHV infection influences the production of multiple cytokines in our *ex vivo* infection model, (2) lower cytokine levels and/or repression of cytokines during infection are generally associated with higher susceptibility to KSHV infection, (3) several individual cytokines show significant negative associations with susceptibility to KSHV infection and (4) IL-21 is positively correlated with KSHV infection of plasma cells. Overall, these data suggest that distinct inflammatory responses in each tonsil specimen contribute to variable susceptibility to KSHV infection.

### IL-21 supplementation increases KSHV infection in tonsil B lymphocytes

Because IL-21 production was positively correlated with plasma cell infection in our initial dataset (Fig 1E), we wanted to examine the impact of manipulating IL-21 levels on the establishment of KSHV infection. To do this, we performed Mock infection or KSHV infection in 12 unique tonsil samples and supplemented the resulting cultures with varying concentrations of recombinant IL-21. At 3 dpi, we analyzed these cultures for GFP+ B lymphocytes by flow cytometry to assess the magnitude of KSHV infection (Fig 2A & B). Although the specimens included in this data set had high variability in their baseline susceptibility, we can see increased infection in response to IL-21 treatment, and the effect seems to be particularly strong in the more susceptible samples (Fig 2A). Normalization of the data to each specimen’s untreated control reveals that at 10/12 samples show increased infection upon treatment with 100pg/ml of IL-21. Importantly, most of these concentrations were higher than what was observed by 3 dpi in our initial dataset quantitating native cytokine secretion in our culture system (Fig 1A), which may explain why we didn’t observe an association of IL-21 secretion with overall infection at that timepoint (Fig 1D). We then repeated these supplementation experiments with only the 100pg/ml dose of recombinant IL-21 in an additional 12 tonsil specimens and examined both overall infection and subset-specific responses in these cultures at 3 dpi using our B cell immunophenotyping panel (Table 1 and Supplemental Fig 1A). Similar to the initial dataset, this analysis shows increased infection in response to recombinant IL-21 in the majority of tonsils, and the difference in GFP+ B lymphocytes was statistically significant in IL-21 treatment compared to control (p=0.02, F=6.4) (Fig 2C). This increase in infection was not associated with alterations in the frequency of viable B cells in the cultures (Supplemental Table 1B).

**Figure 2:**
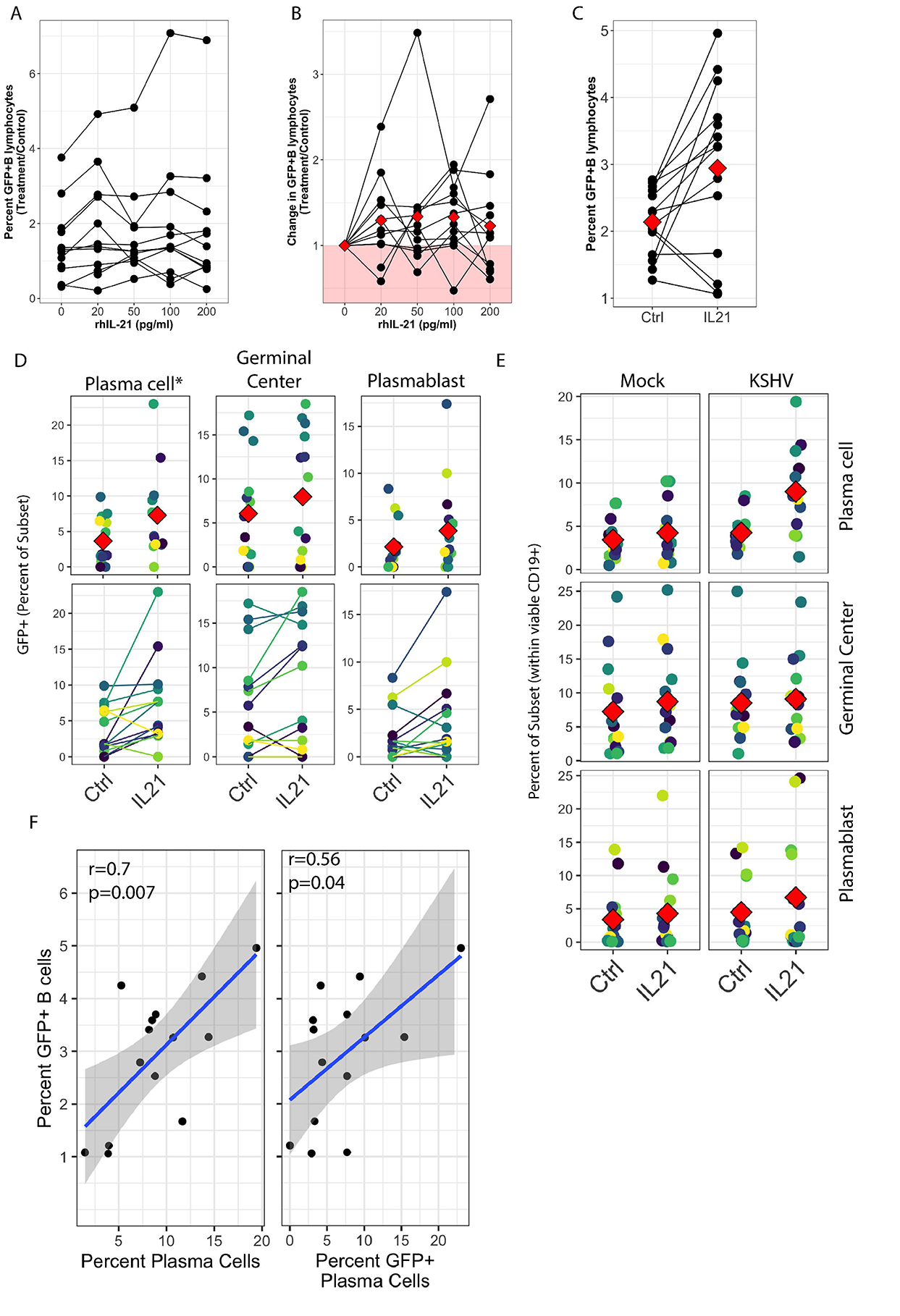
IL-21 supplementation increases overall KSHV infection and plasma cell frequencies. Naïve B Lymphocytes from 12 tonsil donors were infected with KSHV.219 and cultured with indicated doses of recombinant human IL-21 and analyzed at 3dpi by flow cytometry (A) the dose effect of IL-21 supplementation on GFP+ viable B lymphocytes. (B) data as in (A) normalized to the untreated control for each specimen. (C) 12 additional tonsil donors analyzed as in (A) with only 100pg/ml IL-21 treatment. Red diamonds indicate group means. p=0.02, F=6.4 via one-way repeated measures ANOVA. Tonsil lymphocyte specimens from (C) were stained for B cell immunophenotypes and analyzed by flow cytometry and GFP for KSHV infection to determine (D) GFP frequencies within B cell subsets for KSHV-infected cultures and (E) total B cell subset frequencies for each condition. Top panels in (D) and (E) show individual sample quantities and means (red diamonds) and bottom panel in (D) shows trends of increased/decreased subset targeting on a per-sample basis. Colored points denote unique tonsil specimens and can be compared between panels D and E. See Supplemental Table 1A for full statistics for all subsets for panel (D) and Supplemental Table 1B for full statistics for all subsets in panels (C) and (E). (F) Pearson correlation between overall GFP+ B cells in KSHV-infected, IL-21 treated cultures as in (C) and the level of plasma cells (right) and infection of plasma cells (left). Blue line is linear model regression and grey shading indicates 95% confidence interval.

**Table 1:**
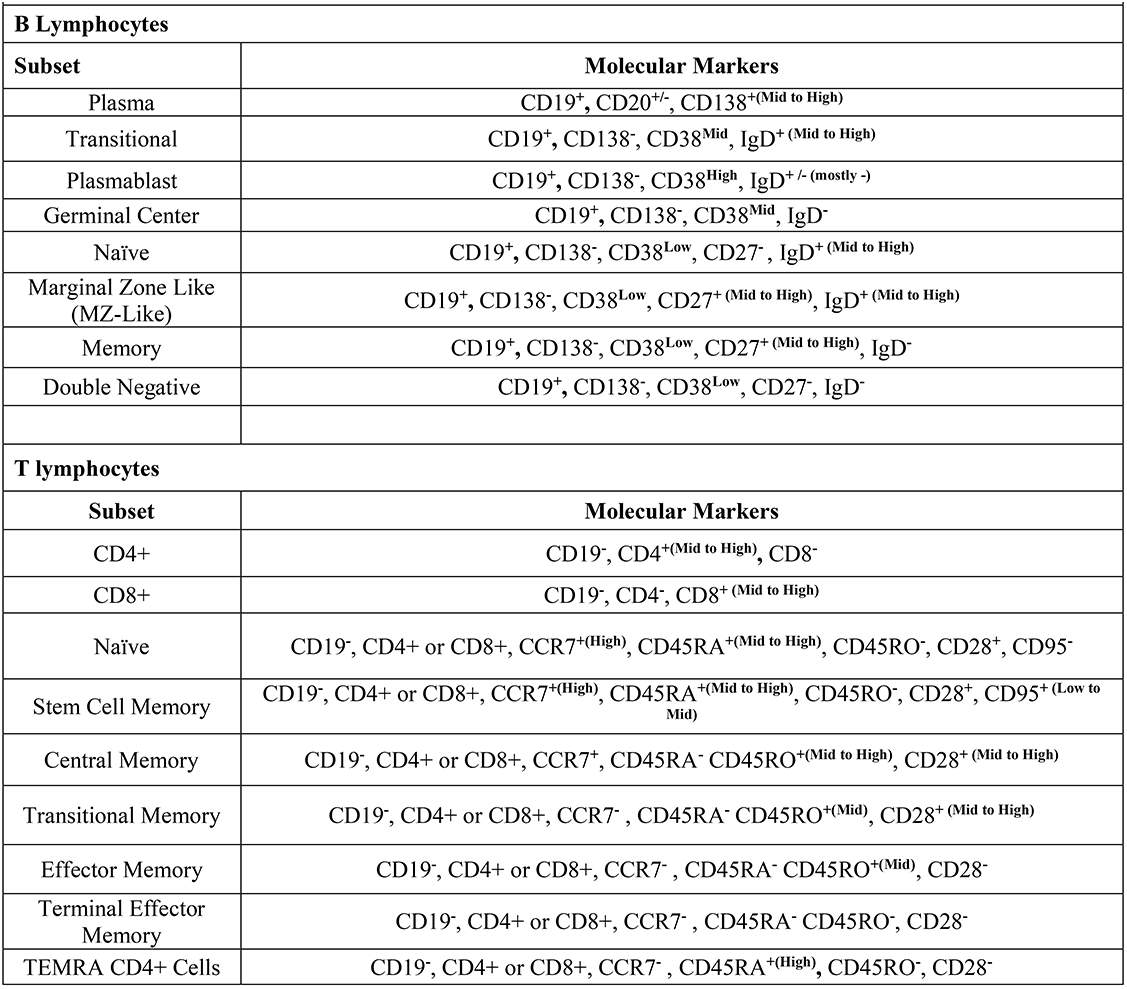
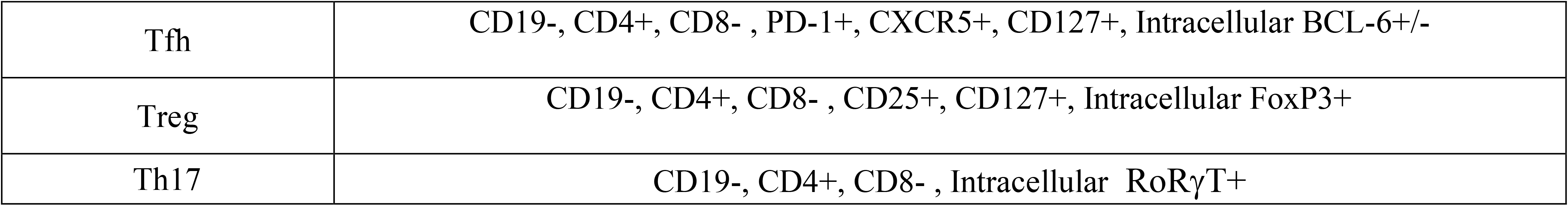
Lineage definitions for lymphocyte subsets used in the study.

### IL-21 increases plasma cell frequency and susceptibility in primary human tonsil B lymphocytes

To examine whether IL-21 treatment altered the B cell subset-specific distribution of KSHV infection in these experiments, we quantitated the percent of each B cell subset that was GFP+ to determine the within-subsets distribution of KSHV infection in control or IL-21 treated cultures. One-way repeated measures ANOVA analysis indicates that IL-21 supplementation did not significantly affect KSHV infection for most subsets (Supplemental Table 1A). However, we observed a significant increase in plasma cell targeting with IL-21 treatment (p=0.02, F=6.6) and notable, but non-significant, increases in the infection of germinal center and plasmablast subsets (Fig 2D). We next examined whether IL-21 supplementation was associated with alterations in B cell frequencies in either Mock or KSHV-infected cultures. Two-way repeated measures ANOVA analysis (Supplemental Table 1B) revealed a highly significant increase in total plasma cell frequency associated with both IL-21 treatment and KSHV infection with a significant interaction of the two variables (Fig 2E). Neither infection nor treatment had a significant effect on germinal center cell frequencies, but there was a significant main effect of KSHV infection on frequencies of plasmablasts in these cultures (p=0.03, F=6). Post hoc paired T tests revealed significant differences with IL-21 treatment on total plasma cells (p=0.0002) for the KSHV-infected conditions only, and in IL-21 treated conditions there was a significant difference between Mock and KSHV cultures for total plasma cells (p=0.0003). In order to determine whether the increase in total plasma cell frequency and/or increased infection of plasma cells was directly correlated with the effect of IL-21 on total KSHV infection, we performed linear model regressions and analyzed the results using Pearson’s method (Fig 2F). These results reveal a significant linear correlation between total GFP and plasma cell frequency (r=0.7, p=0.007) and a weaker, but still significant, correlation between total GFP and the frequency of GFP+ cells within the plasma cell subset (r=0.56, p=0.04). Taken together this data shows that IL-21 treatment promotes the establishment of KSHV infection in human tonsil lymphocytes, and that this increased infection is correlated with both increased plasma cell frequencies and increased plasma cell infection at 3 dpi.

### Neutralization of IL-21 inhibits KSHV infection in primary tonsil B lymphocytes

We next wanted to determine whether neutralization of the natively-secreted IL-21 in our tonsil lymphocyte cultures would affect the establishment of KSHV infection. To do this, we performed infections with Mock or KSHV-infection in 11 unique tonsil specimens, included varying concentrations of an IL-21 neutralizing antibody in the resulting cultures, and assessed the magnitude and distribution of KSHV infection at 3 dpi by flow cytometry. These results reveal decreased KSHV infection in the presence of IL-21 neutralizing antibodies (Fig 3A). One-way repeated measures ANOVA revealed a significant effect of IL-21 neutralization on GFP+ cells in KSHV infected cultures (p=0.00001, F=9.4) and post-hoc Dunnett test revealed significance at the 100μg/ml dose (p=0.03). When each sample was normalized to its untreated control, we observed that 9/11 samples had decreased infection in the presence of 100μg/ml IL-21 neutralizing antibody and this increased to 10/11 samples at higher antibody doses (Fig 3B).

**Figure 3:**
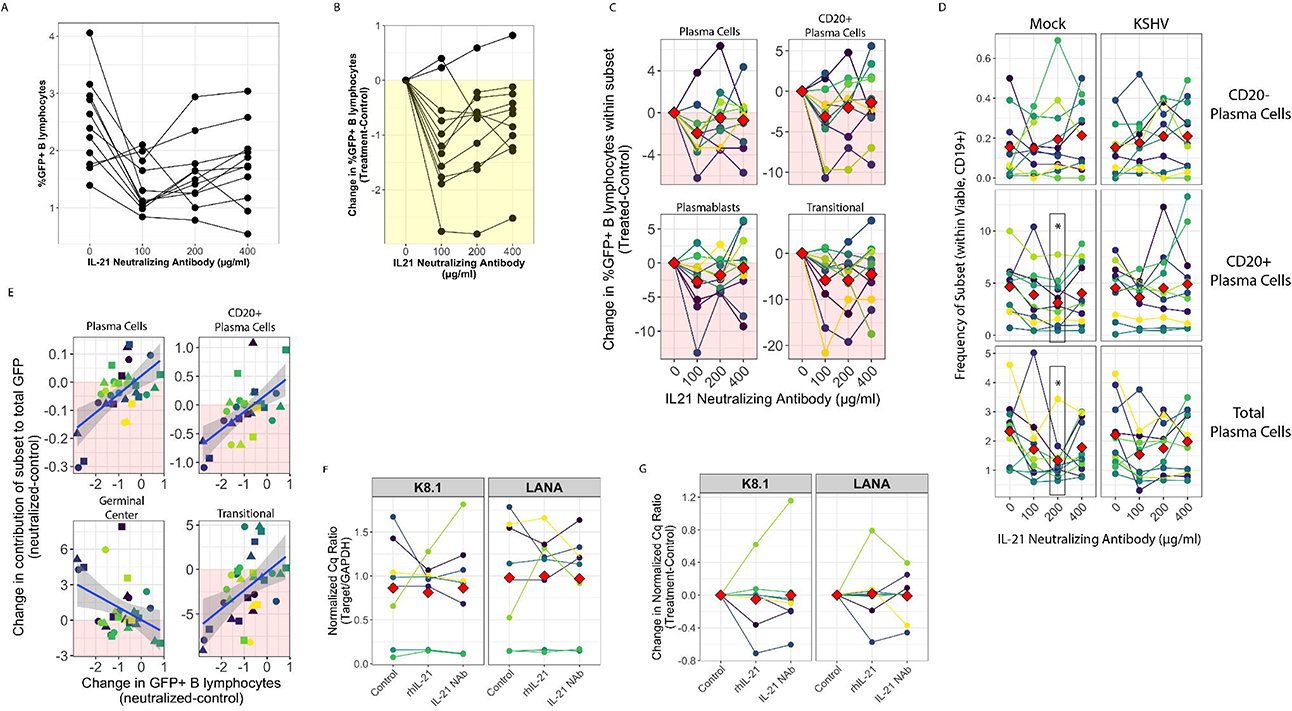
IL-21 neutralization inhibits the establishment of KSHV infection. Naïve B cells from 11 unique tonsil specimens were Mock or KSHV-infected and indicated concentrations of IL-21 neutralizing antibody was added to the resulting total lymphocyte cultures. Cultures were analyzed at 3 dpi for B lymphocyte immunophenotypes and the distribution of KSHV infection via GFP expression. Total GFP+ viable B lymphocytes represented as (A) raw percentages or (B) normalized to the untreated control for each tonsil sample. For (A) one-way repeated measures ANOVA shows p=0.00001, F=9.4 for the main effect of IL-21 neutralization and Dunnett’s test reveals p=0.03 at the 100μg/ml dose. (C) Effect of indicated doses of IL-21 neutralizing antibody on KSHV infection of indicated B cell subsets normalized to the untreated frequency of each subset within each tonsil sample. One-way repeated measures ANOVA on the raw data reveals a significant effect on infection of transitional B cells (p=0.01, F=4.0). Full statistical analysis for all subsets can be found in Supplemental Table 2A. (D) frequencies of plasma cell subsets in the cultures. Full statistical analysis for all subsets can be found in Supplemental Table 2B. Post-hoc paired T-tests showed significant effect of 200μg/ml neutralizing antibody on total plasma cells (p=0.02) and CD20+ plasma cells (p=0.005) in Mock cultures only. (E) Correlation between the effect of IL-21 neutralization on overall infection (x-axis) and the effect of IL-21 neutralization on the contribution of indicated subsets to KSHV infection (y-axis). Shapes indicate doses in this panel (circle=100μg/ml, triangle=200μg/ml, square=400μg/ml). Statistics from Pearson’s linear correlation are as follows: CD20+ plasma cells (r=0.6, p=0.0002), plasma cells (r=0.6, p=0.0003), transitional (r=0.5, p=0.004), germinal center (r=−0.4, p=0.02). For panels C-E colors indicate unique tonsil specimens and can be compared between these panels and red diamonds indicate the mean value for all tonsil specimens. RT-PCR analysis of KSHV transcripts at 3 dpi in 8 unique tonsil specimens with either IL-21 supplementation at 100 pg/ml (Fig 2) or IL-21 neutralizing antibody at 100μg/ml with LANA (latent) and K8.1 (lytic) transcript targets. No RT controls were used to determine that RT-PCR signal is not due to DNA contamination (F) Cq values for viral targets normalized to the within-sample Cq for GAPDH (G) GAPDH-normalized values further normalized to the within-sample value for the untreated control.

Interestingly, when we examined whether this decrease in KSHV infection was associated with alterations in infection of any particular B cell subsets by one-way repeated measures ANOVA (Supplemental Table 2A), we only observed a significant decrease in GFP+ transitional B cells (p=0.01, F=4.0). However, there are non-significant trends showing lower frequencies of infection within plasmablast and CD20+ plasma cell subsets in most samples. (Fig 3C). Moreover, there were no KSHV-specific effects of IL-21 neutralization on the total frequency of any B cell subsets in these experiments via two-way repeated measures ANOVA (Supplemental Table 2B). However, total and CD20+ plasma cell frequencies were significantly reduced at the 200μg dose only in mock cultures (Fig 3D). The observation that the effect of IL-21 neutralization on plasma cell frequencies is restricted to mock-infected cultures is interesting in the context of our IL-21 supplementation data where we observed significant main effects of both IL-21 and KSHV infection on plasma cell frequencies as well as a significant interaction between the two factors (Fig 2E & Supplemental Table 1B). The two data sets taken together support several interesting conclusions: (1) IL-21 affects plasma cell frequencies independent of KSHV infection; evidenced by opposite significant effects of IL-21 supplementation and neutralization in mock cultures, (2) KSHV infection affects plasma cell frequencies independent of IL-21 signaling; evidenced by significant main effect of infection in supplemented cultures and a lack of inhibition in KSHV-infected neutralized cultures, and (3) IL-21 and KSHV can synergistically affect plasma cell frequencies; evidenced by the significant interaction effect and the significant increase in plasma cell frequencies in KSHV-infected cultures that are supplemented with IL-21.

We believe it is the sample-specific variability in responses to IL-21 neutralization that resulted in non-significant effects when the data was parsed to infection within subsets. This variability is not surprising given that magnitude of any effect of neutralization is dependent upon the quantity of IL-21 signaling in the particular culture, which is variable depending upon the sample (Fig. 1A) We hypothesized that if IL-21 signaling is affecting overall infection by contributing to differentiation of KSHV-infected cells, subsets whose differentiation is important to the establishment of infection would accumulate within the GFP+ population with IL-21 neutralization, and this accumulation would correlate with decreased overall levels of infection in response to neutralization. Conversely, subsets whose targeting promotes overall infection in an IL-21 dependent manner would be less represented within the GFP+ population in samples where neutralization was effective at reducing total GFP+ cells. Thus, we calculated the change in total GFP+ B cells in IL-21 neutralized cultures compared to matched control cultures (neutralized-control; effect of neutralization on total infection) and compared this to the change (neutralized-control; effect of neutralization on frequency of subset within GFP+) in the contribution of each subset to infection (between subsets frequency of GFP calculated as GFP+ within subset * frequency of subset within viable CD19+), and performed correlation analysis using Pearson’s method. This analysis reveals that decreased overall infection with IL-21 neutralization was significantly correlated with a decreased contribution of plasma cells (r=0.6, p=0.0002), CD20+ plasma cells (r=0.6, p=0.0002), and transitional B cells (r=0.5, p=0.003) and an increased proportion of infected germinal center cells (r=-0.4, p=0.02) (Fig 3E). As expected, based on the variable per-sample responses we observed for neutralization (Fig 3A&B), these correlations were driven more by sample-specific differences (indicated by point color) than neutralizing antibody dose (indicated by point shape). This data could indicate that IL-21 signaling increases the overall establishment of KSHV infection in tonsil lymphocytes by driving differentiation of germinal center cells into transitional and CD20+ plasma cells. Our previous studies demonstrated that plasma cells display a mixture of lytic and latent KSHV infection [10]. Therefore, we wanted to determine whether the increase in overall KSHV infection with IL-21 treatment and decrease in infection with IL-21 neutralization is due to IL-21-mediated alterations in KSHV lytic reactivation. To examine this, we performed RT-PCR for LANA (latent) and K8.1 (lytic) on total RNA from untreated, IL-21 supplemented or IL-21 neutralizing antibody treated, KSHV-infected cultures from 8 unique tonsil specimens. GAPDH was used as a housekeeping gene and normalizing factor for the viral gene expression data. This data is consistent with our previous data showing a mix of lytic and latent transcripts in infected lymphocytes [10]. These data reveal no significant influence of either supplementation or neutralization on lytic gene expression (Fig 3F). In the majority of samples K8.1 expression remained unchanged or changes were also reflected in LANA transcripts, indicating higher overall infection rather than increased lytic activity (Fig 3G).

### Baseline frequencies of IL21 receptor expression in B cell subsets correlate with susceptibility to KSHV infection

Our data presented thus far demonstrates that IL-21 signaling has a positive effect on the overall establishment of KSHV infection (Fig 2C and 3B) and this increase in overall infection is related to both the frequency of plasma cells (Fig 2E&F and 3D), and the establishment of infection in plasma cells (Fig 2D&F and 3E). Moreover, our data suggests that the increase in plasma cell numbers and targeting may be due to differentiation of new plasma cells via a process that requires IL-21 signaling to germinal center B cells (Fig 3E). To further address the early stages of the IL-21 response during infection, we examined expression of the IL21 receptor in primary human tonsil B lymphocytes at baseline (day 0) in each tonsil specimen. We observed that IL-21 receptor expression is rare on B cells in tonsil at less than 3% of total viable B cells in most samples (Fig 4A). The distribution of IL-21 receptor positive cells among B cell sub-populations is broad and varies substantially between samples, but IL-21 receptor-expressing B cells are most likely to have an MZ-like or plasmablast immunophenotype (Fig 4B) and on a per-sample basis, either plasmablast or MZ-like subsets dominated the IL-21R positive cells in most tonsil samples (Fig 4C). When we examined subset-specific levels of IL-21R via the geometric mean fluorescence intensity of the IL-21R staining within IL-21R+ cells in each subset, we observed that naïve, MZ-like and memory subsets (classical memory and double negative) expressed the highest levels of IL-21R and thus may be most responsive to IL-21 signaling (Fig 4D).

**Figure 4:**
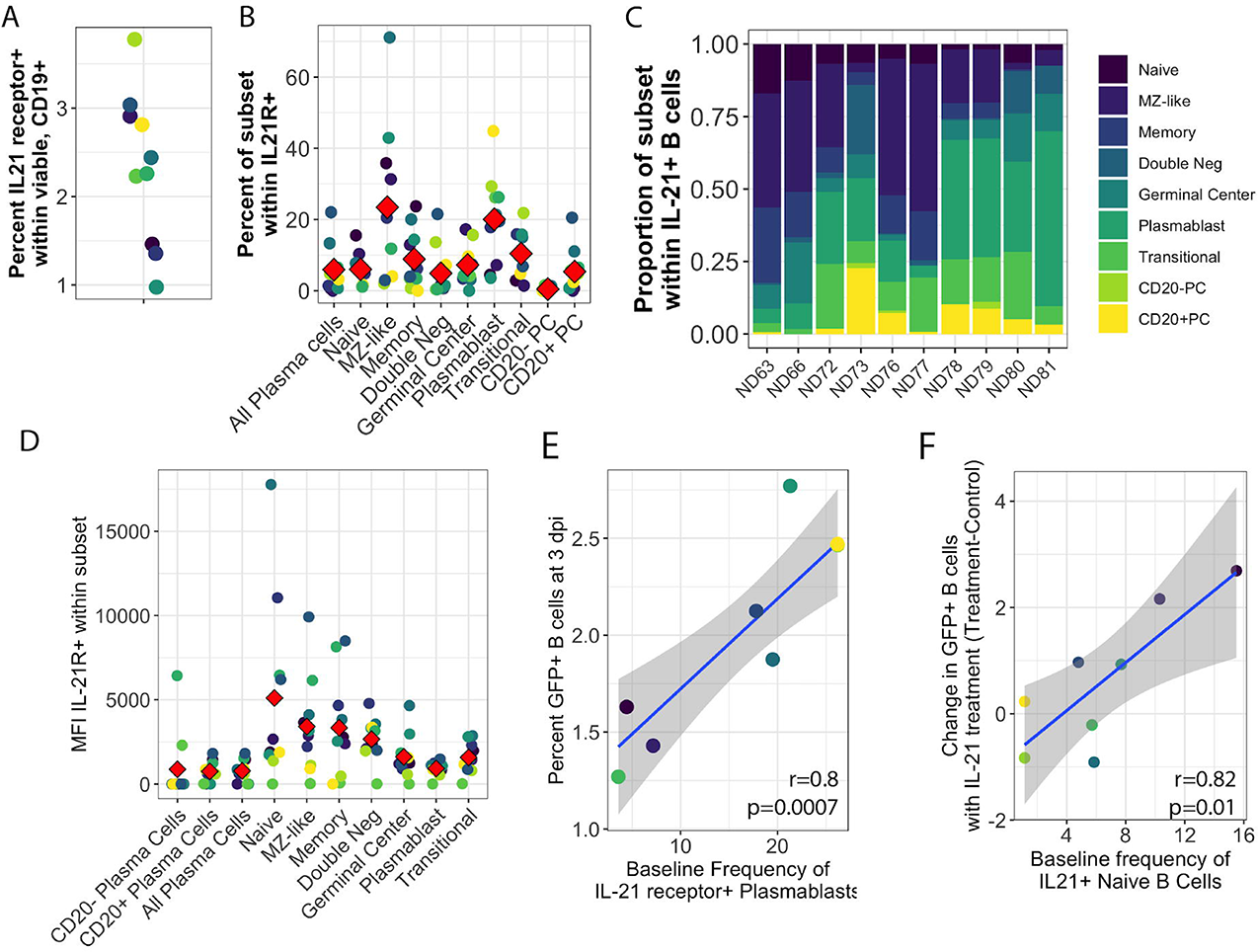
IL21 receptor distribution in primary human tonsil B lymphocytes and its effect on KSHV infection and the response to IL-21 supplementation. B cell immunophenotyping analysis including IL-21R was performed at baseline (Day 0) for 10 unique tonsil specimens (Supplemental Figure 1A). (A) total percentage of IL-21R+ within viable CD19+ B cells. (B) Percent of individual B cell subsets within IL-21+ B cells. Red diamonds indicate the mean value for all tonsil specimens and (C) distribution of B cell subsets within IL-21+ on a per-tonsil basis. (D) MFI of IL-21 receptor within B cell subsets. Red diamonds indicate the mean values for each subset and colors indicate specific tonsil specimens and can be compared between panels (B, D, E and F). (E) Pearson correlation analysis of baseline IL-21R+ plasmablasts with total GFP+ B cells at 3 dpi. Full correlation analysis for all subsets can be found in Supplemental Table 3A (F) Pearson correlation analysis of baseline IL-21R+ naïve B cells with the effect of IL-21 supplementation on overall KSHV infection in the same tonsil specimens at 3dpi. Full correlation analysis for all subsets can be found in Supplemental Table 3B.

In order to determine whether IL-21 receptor expression prior to infection influenced the establishment of KSHV infection, we aggregated the untreated conditions from both the supplementation and the neutralization experiments and examined correlations between baseline IL21R distribution and KSHV infection based on overall GFP (Supplemental Table 3A). This analysis revealed that the proportion of plasmablasts within IL21R+ B cells is significantly correlated with overall susceptibility to KSHV infection in the absence of any IL-21 treatment (r=0.81, p=0.0007) (Fig 4E). However, in experiments where we supplemented cultures with IL-21, the IL-21-mediated increase in overall KSHV infection at 3 dpi (Fig 2C) is correlated with the baseline frequency of IL-21 receptor expression on naïve B cells (r=0.82, p=0.01) (Fig 4F and Supplemental Table 3B). However, there were no significant correlations between the MFI of IL-21R at baseline and total GFP at 3 dpi for any subset (Supplemental Table 4D). These results suggest that IL-21+ plasmablasts are important for susceptibility to KSHV in the absence of high levels of IL-21 at early timepoints in untreated cultures while naïve B cells contribute to the effect of IL-21 supplementation.

Interestingly, there was no positive correlation seen between baseline IL-21R expression and the response of plasma cell frequencies to IL-21 (Supplemental Table 3C), suggesting that the plasma cell response to IL-21 at 3 dpi in KSHV infected cultures may be a product of IL-21 receptor up-regulation in response to infection instead of intrinsic baseline levels of IL-21 on plasma cells or plasma cell precursors in our tonsil lymphocyte cultures. Indeed, modulation of IL-21 receptor expression by KSHV infection is one possible mechanism for the synergistic promotion of plasma cell numbers we observe with both IL-21 treatment and infection (Fig 2D).

### IL-21R+ plasmablasts increase in response to KSHV infection and IL-21R+ Plasma cells increase in response to IL-21 only in KSHV+ cultures

In order to examine this hypothesis, we analyzed IL-21R expression on B cell subsets at 3 dpi in our culture system with or without IL-21 supplementation to determine whether KSHV and/or IL-21 can modulate the response to IL-21 during infection. There were no statistically significant differences in either the frequency (Fig 5A) or fluorescence intensity (Fig 5B) of IL-21R with KSHV infection or IL-21 supplementation. When we examined IL-21R expression on KSHV-infected (GFP+) cells vs GFP-cells in the same culture, we observed non-significant trends in the data showing that GFP+ cells were more likely to be IL-21R+ (Fig 5C) and had higher MFI for IL-21R expression (Fig 5D) compared to GFP-cells. Neither of these effects was altered by IL-21 stimulation. When we examined the distribution of B cell subsets within IL-21R+ B cells, the majority of effects on IL-21R expression at 3 dpi were present in both Mock and KSHV-infected cultures, indicating they are a product of the culture system and not driven by KSHV (Supplemental Tables 4A&B). However, the proportion of plasmablasts within IL21R+ cells was significantly increased comparing Mock to KSHV-infected cultures without IL-21 treatment (p=0.03) and this difference was further increased with the combination of KSHV infection and IL-21 treatment (Fig 5E and Supplemental Table 4B). Comparing untreated and IL-21 treated cultures within infection conditions revealed a significant effect of IL-21 treatment on IL-21+ plasma cells only in the KSHV-infected cultures (Fig 5E and Supplemental Table 4C). However, there was no significant effect of KSHV infection or IL-21 treatment on the fluorescence intensity of IL-21R for any subset (Supplemental Table 4D). Interestingly, there was a significant correlation between the frequency of plasmablasts within IL-21+ and overall infection at 3dpi (Fig 5F, left), which was driven by samples where large increases in GFP in response to IL-21 treatment corresponded to large increases in IL-21R+ plasmablasts (Fig 5F, right). This result is particularly interesting taken together with the correlation between baseline plasmablast frequencies and overall infection at 3 dpi in untreated conditions.

**Figure 5:**
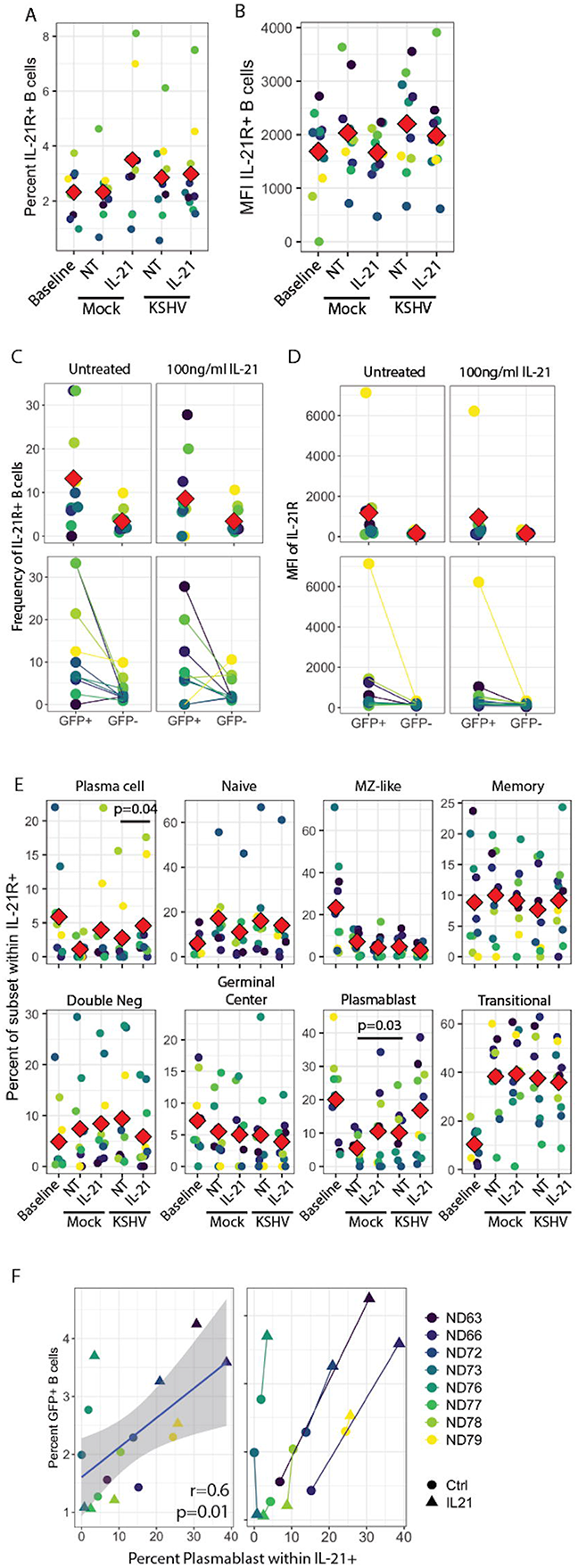
IL-21R+ plasmablasts increase in response to KSHV infection and IL-21R+ Plasma cells increase in response to IL-21 only in KSHV+ cultures. (A) Total percentage of IL-21R+ B cells at baseline and 3dpi within Mock, Mock+100pg/ml IL-21, KSHV, KSHV+ 100pg/ml IL-21 conditions. (B) conditions as in (A) for mean fluorescence intensity of IL-21R staining in IL-21R+ B cells (C) Frequency and (D) MFI of IL-21R for IL-21R+ B cell within untreated GFP+, GFP-and 100pg/ml IL-21 GFP+, GFP-cells. Top panels in (C) and (D) show individual sample quantities and means (red diamonds) and bottom panels show trends of increase/decrease comparing GFP+ to GFP-within the same culture. (E) Distribution of IL-21 receptor on B cell subsets at day 0 (baseline) or 3dpi within Mock, Mock+100pg/ml IL-21, KSHV or KSHV+ 100pg/ml IL-21 conditions. Red diamonds indicate the mean values for each condition and significant differences were assessed via two-way repeated measures ANOVA (Supplemental Table 4A) and post-hoc paired T-tests for both culture/infection conditions (Supplemental Table 4B) and IL-21 treatment (Supplemental Table 4C). (F) Pearson correlation analysis of the frequency of plasmablasts within IL-21+ and overall infection at 3dpi (left). GFP response (y=axis) and plasmablast response x-axis to IL-21 treatment for each sample (right).

These results may indicate that infection and IL-21 treatment is affecting IL21R expression on existing plasmablasts and plasma cells, or that KSHV and IL-21 synergistically drive differentiation of IL-21R+ cells to plasmablast and plasma cell phenotypes. Our observation that IL-21R+ naïve B cells at day 0 are correlated with the response of KSHV infection to IL-21 treatment (Fig 4F) is one indication that differentiation may be playing a role in the IL-21 response. However, our data do not exclude the possibility that a combination of both receptor modulation and differentiation are contributing to the observed 3 dpi phenotypes in the presence of both IL-21 and KSHV infection.

### Characterization of T cell subsets producing IL-21 in primary human tonsil B lymphocytes

We next wanted to determine the source of native IL-21 secretion in our culture system, and determine whether the production of IL-21 is affected by KSHV infection. To accomplish this, we utilized an additional immunophenotyping panel for T cell subsets (Table 2 and Supplemental Fig 1B), and performed intracellular cytokine staining (ICCS) on unstimulated T cells at 3 dpi to identify T cell subsets that are producing IL-21 (Supplemental Fig 1C) in Mock and KHSV-infected cultures from 14 unique tonsil samples. This data shows that IL-21 secretion in T cells is highly variable between tonsil lymphocyte cultures (range=1.4-24.7%, mean=10.6, median=10.55, standard deviation=6.9), but is not significantly affected by KSHV infection (Fig 6A). More of the IL-21+ cells were CD4+ T cells vs. CD8+ T cells and this distribution was also not affected by KSHV infection (Fig 6B). We next utilized a metric called integrated MFI (iMFI) [27] to examine the contribution of T cell subsets to IL-21 secretion. This value is calculated by multiplying the mean fluorescence intensity of IL-21 in each T cell subset by the subset’s frequency within total T cells. Thus, iMFI integrates the amount of IL-21 being secreted by a subset with the frequency of that subset; more correctly quantitating the contribution of low frequency subsets that are high IL-21 producers. Unlike the frequency of IL-21+ T cells, the iMFI of IL-21 within CD4+ and CD8+ T cells displayed some overlap (Fig 6C), indicating that, although they are low frequency, CD8+ T cells can be high producers of IL-21. Indeed, there were two samples in our analysis where, based on iMFI, CD4+ and CD8+ T cells were contributing equally to total IL-21 secretion (Fig 6D, red box). When we examined the subset-level distribution of IL-21 secretion within T cells, we observed that within CD4+ cells CD45RO+, central memory, Tfh and RoRγT+ cells displayed the highest frequency of IL-21 positive cells. Among CD8+ T cells, CD45RA+, stem cell memory, central memory and RoRγT+ cells had the highest frequency of IL-21 positive cells (Fig 6E). When we examined whether KSHV infection altered the frequency of IL-21+ T cell subsets, we found that the frequency of IL-21+ CD4+ CD45RA+, CD4+ RoRγT+ and CD8+ central memory subsets was significantly decreased in KSHV-infected cultures vs. Mock cultures. Importantly, KSHV infection did not significantly change the overall levels and subset distribution of T cells within these cultures, indicating these are biological changes within T cell subsets and not due to changes in the T cell population during infection (Supplemental Fig 2A). When we performed the iMFI calculation on the subset level we observed that, as expected based on their established function, CD4+ Tfh had an increased contribution to IL-21 secretion relative to their frequency. CD45RO+ T cells contributed more to IL-21 in CD4+ T cells whereas CD45RA+ T cells contributed more to IL-21 secretion in the CD8+ population. Interestingly, among the CD8+ T cell subsets, RoRγT+ and central memory subsets showed increased contribution to IL-21 secretion relative to their frequency indicating that these cells are high IL-21 producers (Fig 6F). Finally, KSHV infection significantly decreased the iMFI of CD4+ CD45RA+ and CD4+ RoRγT+ T cell subsets, which is likely related to their significantly decreased frequency (Fig 6E) rather than an effect on the level of IL-21 secretion from the subset.

**Figure 6:**
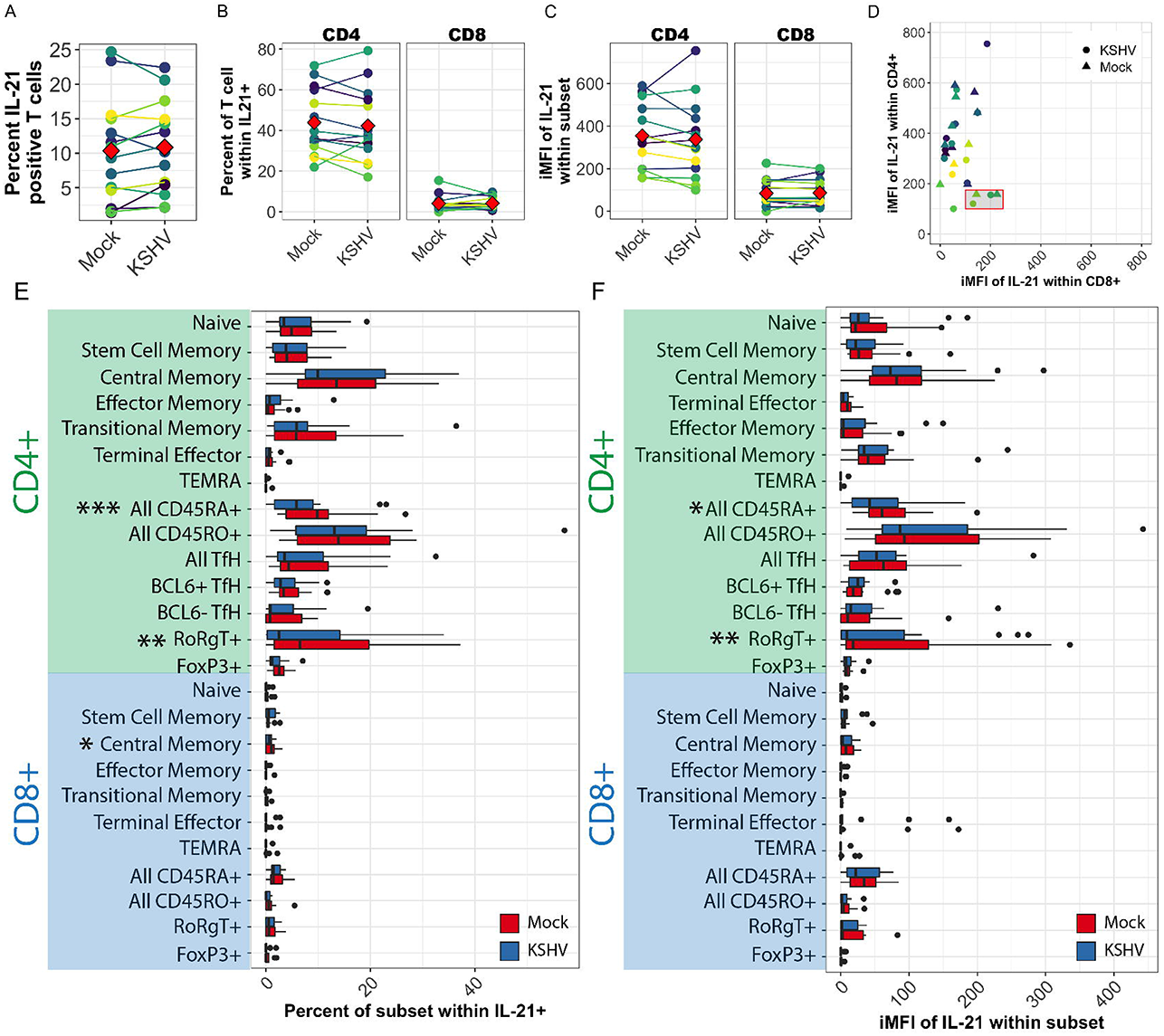
Characterization of T cell subsets producing IL-21 in primary human tonsil B lymphocytes. T cells were analyzed by surface immunophenotyping, intracellular transcription factor staining and ICCS for IL-21 secretion (Supplemental Figure 1B&C) in Mock and KSHV-infected total lymphocyte cultures at 3 dpi in 14 unique tonsil specimens. (A) Total IL-21+ viable non-B cells (B) percent of CD4+ or CD8+ T cells within IL-21+ (C) iMFI of IL-21 within CD4+ and CD8+ T cells in Mock and KSHV culture. For (A), (B) and (C) red diamonds indicate the mean value for the condition and colors indicate specific tonsil specimens and can be compared between the panels. (D) iMFI of IL-21 within CD4+ vs CD8+ T cells in Mock and KSHV-infected cultures (indicated by shape). Red box denotes samples where CD4+ and CD8+ iMFI are comparable (E) Frequency of T cell subsets within IL-21+ in Mock (red) and KSHV-infected (blue) cultures *p=0.05; **p=0.04; ***p=0.003 (F) data as in (E) for iMFI of T cell subsets within IL-21+

### KSHV+ plasma cells are associated with a specific combination of IL-21 producing T cells

We wanted to determine whether IL-21 secretion by any particular T cell subset was correlated with susceptibility to KSHV infection in our experiments. To do this, we performed B cell immunophenotyping analysis to determine the extent and distribution of KSHV infection in the same cultures where ICCS was performed on T cell subsets. When we examined correlations between IL-21 secretion by T cell subsets and overall KSHV infection in B cells (Supplemental Table 5A) we found that only the frequency of IL-21+ CD8+ central memory cells were significantly correlated (r=0.57, p=0.03) (Fig 7A) and there were no significant correlations between the iMFI of IL-21 in T cell subsets and overall GFP+ B cells at 3dpi (Supplemental Table 5B). When we examined correlations between baseline T cell subsets and infection at 3dpi, the only significant correlation was a negative impact of CD4+ CD45RO+ T cells (r=-0.57, p=0.03) (Supplemental Table 5C). Given that these experiments rely on native IL-21 secretion over time in the culture system, as opposed to high levels of recombinant IL-21 added at day 0 in our previous experiments (Fig 2), we hypothesized that impacts on total GFP may be absent at this timepoint because KSHV targeting and manipulation of plasma cell frequencies in response to IL-21 precedes the effect on total infection. Thus, we examined correlations between plasma cell infection in the context of subset-specific IL-21 secretion by T cells. In this analysis, we found that (1) the baseline frequency of CD8+ central memory cells (r=0.59, p=0.02), (2) their frequency within IL-21+ at 3dpi (r=0.82, p=0.0003) and (3) their iMFI at 3dpi (r=0.75, p=0.002) significantly correlated with plasma cell targeting (Fig 7B and Supplemental Figure 2 B-D). Although CD8+ central memory cells are a low-frequency subset, they are significant contributors to IL-21 secretion within CD8+ cells based on iMFI (Fig 6F) and they show redundant correlations in our data that indicate they are important for the early establishment of KSHV infection in tonsil lymphocytes. In addition, the frequency of both CD4+ and CD8+ T cells that express RoRγT+ (the Th17/Tc17-defining transcription factor) within IL-21+ T cells were significantly correlated with plasma cell targeting by KSHV (Fig 7C and Supplemental Figure 2B). These correlations were stronger for CD4+ RoRγT+ cells and were coupled with a significant negative correlation with KSHV-infection of naïve T cells (Supplemental Figure 2B). In this data, we noticed that the same two tonsil specimens were driving the positive correlations between plasma cell targeting by KSHV and IL-21 secretion by CD8+ central memory, CD4+ RoRγT+, and CD8+ RoRγT+ T cells (Fig 7B&C). We next correlated the iMFI of these three T cell subsets (CD8+ central memory, CD4+ RoRγT+, and CD8+ RoRγT+; hereafter referred to as “subsets of interest”) with both GFP+ plasma cells and total plasma cells, which were significantly elevated in KSHV-infected conditions in response to IL-21 supplementation (Fig 2E). Consistent with our previous dataset (Fig 2F), the targeting of plasma cells by KSHV is directly correlated with the total frequency of plasma cells (Fig 7D, top left panels). The iMFI of our T cell subsets of interest was significantly correlated with GFP+ plasma cells but was not correlated to total plasma cell numbers. However, the same two tonsil samples driving the previous correlations did show elevated plasma cell frequencies together with elevated T cell iMFI in this data (Fig 7D, red boxes) while the lack of correlation in this data was driven by samples with high plasma cell frequencies where high KSHV targeting of plasma cells was absent and the iMFI for at least one of the T cell subsets was low. This may indicate that other factors can influence total plasma cell numbers, but both total plasma cells and GFP+ plasma cells are only simultaneously elevated when the IL-21 producing T cells are also present. Moreover, this data indicates that all three IL-21 producing T cells are necessary for the combined plasma cell phenotype. Indeed, we observed direct correlations between the iMFI values of all three T cell subsets of interest, further supporting the conclusion that it is actually the combination of factors rather than independent contributions of each T cell subset that is driving the combined increases in plasma cell frequencies and plasma cell targeting in KSHV-infected conditions.

**Figure 7:**
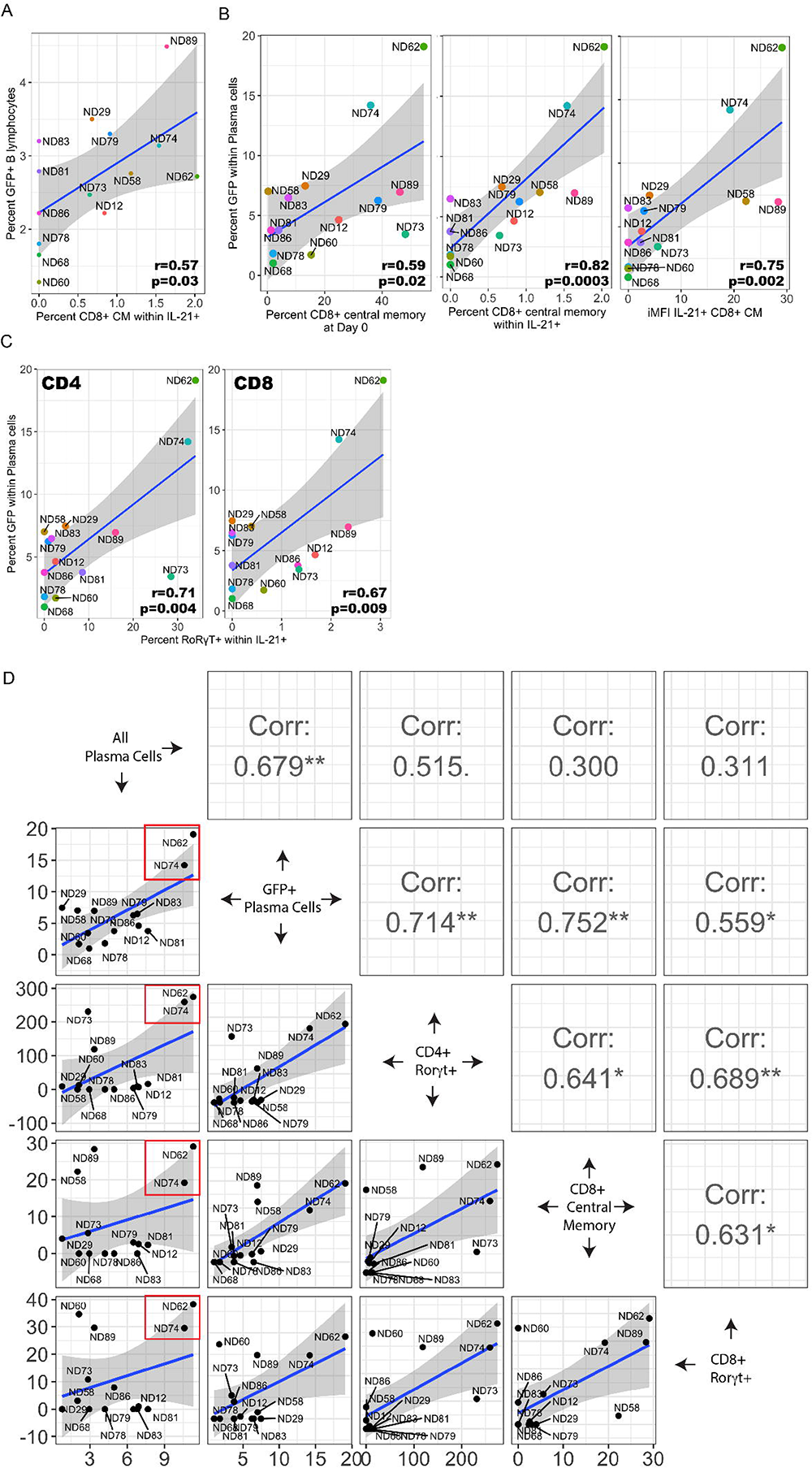
Influence of IL-21 secreting T cell subsets on KSHV infection, plasma cell frequencies and plasma cell targeting. Lymphocyte cultures from the experiments shown in Fig 6 were further analyzed for B cell subsets and the magnitude and distribution of KSHV infection. (A) Pearson correlation analysis of total GFP+ within viable, CD19+ B lymphocytes at 3 dpi with the contribution of CD8+ central memory T cell subsets to IL-21 secretion. Complete statistics for all T cell subsets can be found in Supplemental Table 5A. (B) correlation analysis of GFP+ plasma cells at 3dpi and frequency of CD8+ central memory T cells at day 0 (left), IL-21 secretion by CD8+ central memory T cells at 3dpi (middle), iMFI of IL-21+ CD8+ central memory T cells at 3dpi (right). (C) Pearson correlation analysis of GFP+ plasma cells and frequency of CD4+ (left) or CD8+ (right) RoRγT+ within IL-21+ T cells. (D) pairwise correlation analysis between total plasma cell frequency, GFP+ plasma cell frequency, and iMFI of CD4+ RoRγT+, CD8+ central memory and CD8+ RoRγT+ T cells at 3dpi in KSHV-infected conditions only. Pearson’s correlation coefficients are listed in the top right panels. ***p<0.001, **p<0.01, *p<0.05. For all panels in this figure tonsil sample designations (ND#) are listed adjacent to color-coded data points. Grey shading indicates 95% confidence intervals.

The sample numbers in our dataset were not sufficient for true multivariate analysis with the large number of T cell subsets analyzed. However, multiple pairwise correlations can provide some further insight into this phenomenon. Indeed, we observe highly significant correlations between all three T cell subset based on their frequency at baseline and their frequency at 3dpi (Fig 8A). Interestingly, there were distinctively separate populations of samples in our analysis where the three T cell subsets were either all low or all high, and this distinction was particularly obvious within the 3 dpi frequencies (Fig 8A, right). Similarly, the frequency of these subsets within IL-21+ and the iMFI of IL-21 in these subsets are also significantly correlated, but only in the KSHV-infected samples (Fig 8B). However, the IL-21 correlations were relatively weak compared to the frequency correlations indicating that additional functions of this T cell milieu that are not directly related to IL-21 secretion may also influence the plasma cell targeting phenotype. Finally, we wanted to determine whether any other T cell subsets correlated with all three subsets, and may be additionally contributing to the T cell milieu which promotes plasma cell targeting by KSHV. Since 3 dpi frequency yielded the strongest correlations within the T cell data (Fig 8A, right), we performed pairwise correlations between our three subsets of interest and the remaining T cell subsets in the analysis (Fig 8C). The data reveals that the frequencies of all three subsets of interest are also significantly correlated with the frequency of CD4+ stem cell memory and samples where BCL6+ cells predominate within the CD4+ Tfh subset. Taken together, these results indicate that there is a defined T cell milieu in some tonsil samples that includes elevated frequencies of IL-21-producing CD8+ central memory, CD4+ RoRγT+, and CD8+ RoRγT+ T cells, as well as other T cell subsets, and this particular milieu correlates with the ability of KSHV to increase plasma cell numbers and target plasma cells for infection at early timepoints.

**Figure 8:**
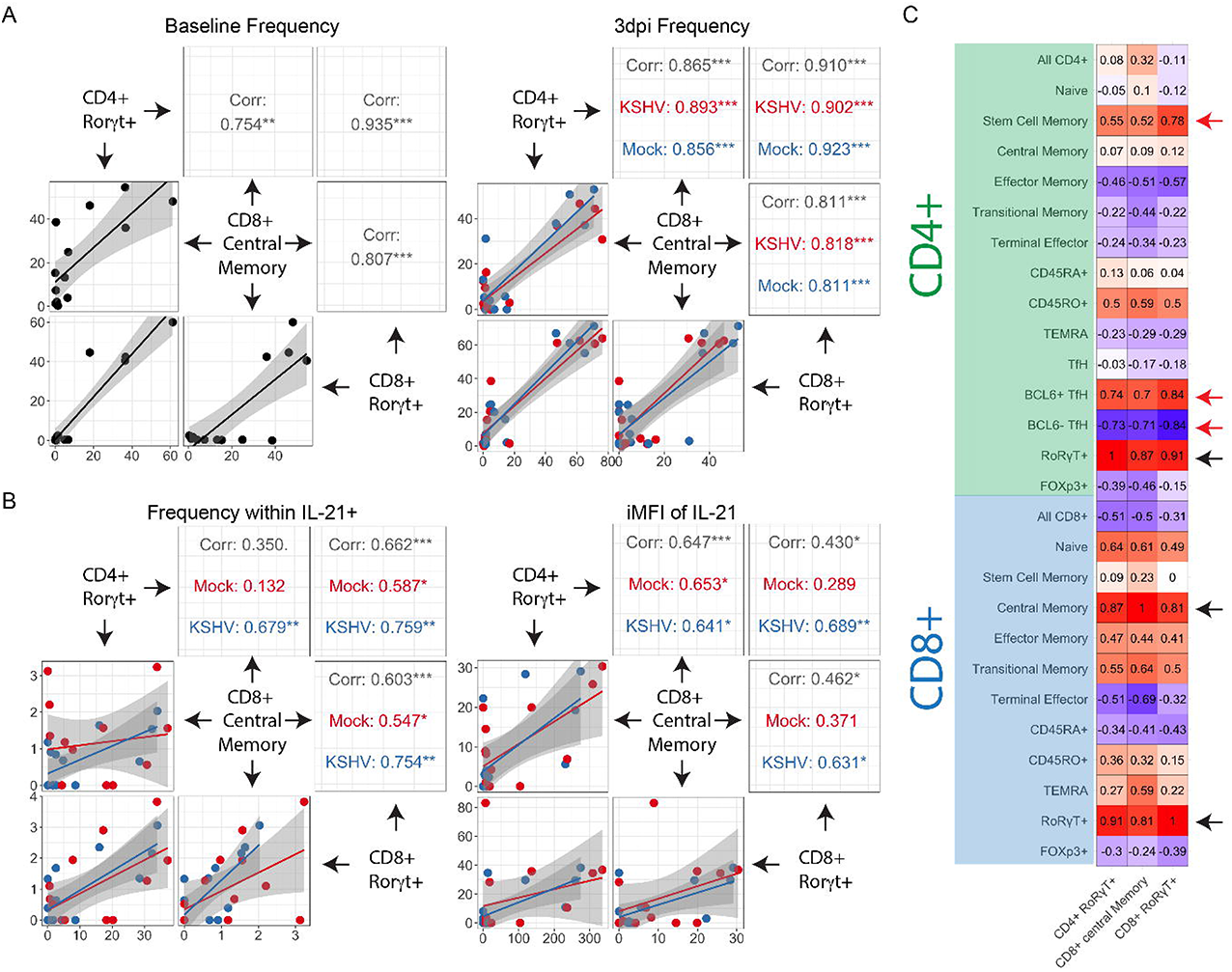
IL-21-producing CD8+ central memory, CD4+ RoRγT+, and CD8+ RoRγT+ T cells indicate a T cell milieu that influences KSHV infection of plasma cells. (A) Pairwise Pearson correlations between T cell subsets of interest (CD4+ RoRγT+, CD8+ central memory and CD8+ RoRγT+) based on their frequency at baseline (left) and their frequency at 3dpi (right) in Mock (blue) or KSHV-infected (red) conditions. **(B)** Pairwise Pearson correlations between T cell subset (CD4+ RoRγT+, CD8+ central memory and CD8+ RoRγT+) based on their frequency within IL-21+ (left) and the iMFI of IL-21 at 3dpi (right) in Mock (blue) or KSHV-infected (red) conditions. For (A) and (B) ***p<0.001, **p<0.01, *p<0.05 and grey shading indicates 95% confidence intervals **(C)** correlogram of pairwise correlations between 3 dpi frequencies of the T cell subsets of interest (CD4+ RoRγT+, CD8+ central memory and CD8+ RoRγT+) (x-axis) and the 3 dpi frequencies of all T cell subsets analyzed (y-axis). Pearson’s r values with an absolute value greater than or equal to 0.53 are statistically significant for this dataset.

## Discussion

Our results presented in this study indicate that KSHV can influence cytokine production in tonsil-derived lymphocytes and that the host inflammatory state contributes to the dramatic variation in susceptibility we observe among our tonsil lymphocyte specimens [10]. This result is not surprising considering dysregulation of the inflammatory environment is a hallmark of all KSHV-associated malignancies [25, 28, 29]. However, the role of the baseline inflammatory environment in the oral cavity as a potentially modifiable susceptibility factor for the acquisition of KSHV infection in humans is an interesting consideration stemming from these results that deserves further study.

In this study, we uncovered a role for IL-21 signaling in the establishment of KSHV infection in tonsil lymphocytes. That IL-21 plays a role in this process is not particularly surprising in the context of the well-characterized role of IL-21 in the closely-related murine MHV-68 model. Specifically, Collins and Speck recently used IL-21R knockout mice to demonstrate that IL-21 signaling is critical for the establishment of MHV68 latency specifically in B cells. Interestingly, this study showed that the mechanisms of decreased infection were related to decreases in both germinal center and plasma cell frequencies as well as decreased infection in both the germinal center and plasma cell compartment at later timepoints post-infection [20], suggesting a critical mechanism for IL-21 in MHV68 transit of the germinal center and differentiation of follicular-derived plasma cells. The current study represents the first examination of similar mechanisms for the importance of IL-21 in primary KSHV infection in human cells. The relationship between plasma cell differentiation and IL-21 is well-characterized in human immunology [14, 15], and we previously showed that plasma cells are highly targeted during early KSHV infection [10]. However, these results are novel and interesting in that they demonstrate direct correlations between plasma cell frequencies, plasma cell infection, and overall susceptibility to KSHV infection (Fig 2F and 7D). Specifically, the synergistic effect of KSHV infection and IL-21 signaling increasing plasma cell frequencies (Fig 2E) plays a role in early infection events that ultimately influences the magnitude of initial dissemination of KSHV within the B cell compartment. Whether this relationship is due a direct effect of the plasma cells themselves or an indirect effect of the process by which IL-21 and KSHV infection manipulates plasma cell frequencies remains to be established, and is the subject of ongoing studies in our laboratory. Moreover, although we have correlative data suggesting that KSHV and IL-21 influence the differentiation of plasma cells (Fig 3E), the data presented herein do not directly interrogate whether KSHV drives B cell differentiation in our tonsil lymphocyte model. Studies are currently ongoing in our laboratory to examine whether KSHV infection influences B cell differentiation and what viral factors influence this process.

Our results herein implicate a particular T cell milieu in promoting plasma cell frequencies and plasma cell targeting during early KSHV infection. Our current analysis identifies IL-21+ CD8+ central memory, IL-21+ CD4+ RoRγT+, and IL-21+ CD8+ RoRγT+ as well as BCL6+ Tfh and CD4+ stem cell memory subsets independent of IL-21 secretion as participants in this milieu. To our knowledge, this particular combination of T cells does not have a previously defined function in tonsillar immunology. It will be interesting to perform true multivariate analysis to establish the contribution of the baseline T cell milieu to KSHV infection once we have analyzed enough unique tonsil specimens to make such analysis feasible. Based upon our current data, we would hypothesize that the T cell composition of tonsil samples at baseline can be used to predict sample-level susceptibility to KSHV infection. Epidemiological evidence from Africa suggests that acquisition of KSHV infection in infants is lower than expected based on shedding of KSHV by household contacts, indicating that unknown factors influence the initial acquisition of KSHV infection in childhood [30]. Our results at least implicate an immunologically activated state in the initial establishment of KSHV infection in tonsil lymphocytes, suggesting that prior pathogen exposure, chronic infection or temporally-associated acute infections may create an inflammatory state in the tonsil that is permissive for KSHV transmission.

Both IL-21 and IL-6, which is highly induced in our KSHV-infected cultures (Fig 1), are involved in the maintenance and function of RoRγT+ T cells via STAT3 signaling. Th17/Tc17 cells produce IL-17A, which is another cytokine that promotes the establishment of chronic MHV68 infection via promotion of the MHV68-mediated germinal center response [31] and is mechanistically linked to suppression of T cell-intrinsic IRF-1 [32]. These results are particularly interesting in light of our current findings showing that IL-21 secretion from, and baseline levels of, RoRγT+ T cells correlate with the early targeting of plasma cells during KSHV infection (Fig 7C and Supplemental Fig 2C), suggesting that the Th17/Tc17 environment in the tonsil may be a critical factor influencing donor-specific susceptibility to KSHV infection. Indeed, as an important site for mucosal immunity in the oral cavity, the Th17/Tc17 environment in tonsil is highly dynamic and physiologically important. In fact, Th17 cells play a major role in host defenses against several pathogens and immunopathogenesis [33, 34]. Many studies have shown that certain parasites modulate the immune response by inducing Th17 [35, 36]. Previous finding suggest that parasite infection is linked with KSHV infection in Uganda [37]. Thus, the parasite burden in sub-Saharan Africa may modulate susceptibility to KSHV infection via manipulating Th17/Tc17 frequencies.

Consistent with the MHV68 literature, our current results mechanistically implicate germinal center cells in these observations (Fig 3E). However, although our *ex vivo* model of KSHV infection in primary lymphocytes is a powerful tool, it certainly does not recapitulate the complex interactions that are needed for a functional germinal center reaction, so further examination of these particular mechanisms will require the utilization of an alternative model system, such as a humanized mouse. However, the participation of CD8+ central memory, Th17 and Tc17 cells in the T cell milieu associated with increased plasma cell targeting indicates that an extrafollicular pathway may also be active in this process, which is consistent with literature implicating extrafollicular maturation of KSHV-infected B cells in the pathogenesis of MCD [38].

Although CD8+ T cells are minor contributors to IL-21 secretion compared to CD4+ T cells, our data strongly indicates that they participate in the inflammatory milieu that promotes KSHV dissemination in our model (Fig 7 and Fig 8). Previous studies have shown that CD8+ T cells can be observed in B lymphocyte areas of tonsil and provide co-stimulatory signals and cytokines to support B cell survival [39]. Interestingly, recent studies have shown that IL-6 regulates IL-21 production in CD8+ T cells in a STAT3-dependent manner, and that CD8+ T cells induced in this way can effectively provide help to B cells [40]. Thus, the induction of human IL-6 during KSHV infection may modulate the function of CD8+ T cells in a way that favors the establishment and dissemination of KSHV infection within the lymphocyte compartment independent of traditional CD4+ helper T cells, which would be an interesting dynamic in the context of CD4+ T cell immunosuppression associated with HIV infection where KSHV-mediated malignancies are common.

## Material and Methods

### Ethics Statement

Human specimens used in this research were de-identified prior to receipt, and thus were not subject to IRB review as human subjects research.

### Reagents and Cell Lines

CDw32 L cells (CRL-10680) were obtained from ATCC and were cultured in DMEM supplemented with 20% FBS (Sigma Aldrich) and Penicililin/Streptomycin/L-glutamine (PSG/Corning). For preparation of feeder cells CDw32 L cells were trypsinized and resuspended in 15 ml of media in a petri dish and irradiated with 45 Gy of X-ray radiation using a Rad-Source (RS200) irradiator. Irradiated cells were then counted and cyropreserved until needed for experiments. Cell-free KSHV.219 virus derived from iSLK cells [39] was a gift from Javier G. Ogembo (City of Hope). Human tonsil specimens were obtained from the National Disease Research Interchange (NDRI; ndriresource.org). Human fibroblasts for viral titering were derived from primary human tonsil tissue and immortalized using HPV E6/E7 lentivirus derived from PA317 LXSN 16E6E7 cells (ATCC CRL-2203). Antibodies for flow cytometry were from BD Biosciences and Biolegend and are detailed below. Recombinant human IL-21 was from Preprotech (200-21) and IL-21 neutralizing antibody was from R&D Systems (991-R2).

### Isolation of primary lymphocytes from human tonsils

De-identified human tonsil specimens were obtained after routine tonsillectomy by NDRI and shipped overnight on wet ice in DMEM+PSG. All specimens were received in the laboratory less than 24 hours post-surgery and were kept at 4°C throughout the collection and transportation process. Lymphocytes were extracted by dissection and maceration of the tissue in RPMI media. Lymphocyte-containing media was passed through a 40μm filter and pelleted at 1500rpm for 5 minutes. RBC were lysed for 5 minutes in sterile RBC lysing solution (0.15M ammonium chloride, 10mM potassium bicarbonate, 0.1M EDTA). After dilution to 50ml with PBS, lymphocytes were counted, and pelleted. Aliquots of 5(10)7 to 1(10)8 cells were resuspended in 1ml of freezing media containing 90% FBS and 10% DMSO and cryopreserved until needed for experiments.

### Infection of primary lymphocytes with KSHV

Lymphocytes were thawed rapidly at 37°C, diluted dropwise to 5ml with RPMI and pelleted. Pellets were resuspended in 1ml RPMI+20%FBS+100μg/ml DNaseI+ Primocin 100μg/ml and allowed to recover in a low-binding 24 well plate for 2 hours at 37°C, 5% CO2. After recovery, total lymphocytes were counted and naïve B cells were isolated using Mojosort Naïve B cell isolation beads (Biolegend 480068) or Naïve B cell Isolation Kit II (Miltenyi 130-091-150) according to manufacturer instructions. Bound cells (non-naïve B and other lymphocytes) were retained and kept at 37°C in RPMI+20% FBS+ Primocin 100μg/ml during the initial infection process. 1(10)^6^ Isolated naïve B cells were infected with iSLK-derived KSHV.219 (dose equivalent to the ID20 at 3dpi on human fibroblasts) or Mock infected in 400ul of total of virus + serum free RPMI in 12×75mm round bottom tubes via spinoculation at 1000rpm for 30 minutes at 4°C followed by incubation at 37°C for an additional 30 minutes. Following infection, cells were plated on irradiated CDW32 feeder cells in a 48 well plate, reserved bound cell fractions were added back to the infected cell cultures, and FBS and Primocin (Invivogen) were added to final concentrations of 20% and 100μg/ml, respectively and recombinant cytokines or neutralizing antibodies were also added at this stage, depending upon the specific experiment. Cultures were incubated at 37°C, 5% CO2 for the duration of the experiment. At 3 days post-infection, cells were harvested for analysis by flow cytometry and supernatants were harvested, clarified by centrifugation for 15 minutes at 15,000 rpm to remove cellular debris, and stored at −80°C for analysis.

### Bead-based immunoassay for supernatant cytokines

Clarified supernatants were thawed on ice and 25μl of each was added to a 13-plex LEGENDplex (Biolegend) bead-based immunoassay containing capture beads for the following analytes: IL-5, IL-13, IL-2, IL-9, IL-10, IL17A, IL-17F, IL-6, IL-21, IL-22, IL-4, TNF-α, and IFN-γ. These assays were performed according to the manufacturer’s instructions, data was acquired for 5000 beads per sample (based on approximately 300 beads per analyte recommended by the manufacturer) using a BD FACS VERSE flow cytometry analyzer and cytokine concentrations in the experimental supernatants was calculated from standard curves using the LEGENDPlex software.

### Flow cytometry analysis of baseline lymphocyte subsets and KSHV infection

Approximately 5(10)^6^ lymphocytes per condition were harvested into a 96-well round bottom plate at day 0 (baseline) or at 3 days post-infection at 1500 rpm for 5 minutes. Cells were resuspended in 100μl PBS containing zombie violet fixable viability stain (BL Cat# 423113) and incubated on ice for 15 minutes. After incubation, cells were pelleted and resuspended in 100ul PBS, containing the following: 2% FBS and 0.5% BSA (FACS Block) was added to the wells. Cells were pelleted at 1500rpm 5 minutes and resuspended in 200ul FACS Block for 10 minutes on ice. Cells were pelleted at 1500rpm for 5 minutes and resuspended in 50μl of PBS with 0.5% BSA and 0.1% Sodium Azide (FACS Wash), **For B cell frequencies** 10μl BD Brilliant Stain Buffer Plus and antibodies as follows: IgD-BUV395 (2.5μl/test BD 563823), CD77-BV510 (2.0 μl/ test BD 563630), CD138-BV650 (2μl/test BD 555462), CD27-BV750 (2μ/test BD 563328), CD19-PerCPCy5.5 (2.0μl/test BD 561295), CD38-APC (10μl/test BD 560158), CD20-APCH7 (2ul/test BL 302313), IgM (2μl/test BL 314524), IgG (2μl/test BD 561298), IgE (2μl/test BD 744319) and IL-21 receptor (2μl/test BD 330114). **For baseline T cell frequencies**. For baseline T cell frequencies 0.5(10)^6^ cells from baseline uninfected total lymphocyte samples were stained and analyzed as above with phenotype antibody panel as follows: CD95-APC (2μl, Biolegend 305611), CCR7-PE (2μl, BD 566742), CD28-PE Cy7 (2μl, Biolegend 302925), CD45RO-FITC (3μl, Biolegend 304204), CD45RA-PerCP Cy5.5 (2μl, 304121), CD4-APC H7 (2μl, BD 560158), CD19-V510 (3μl, BD 562953), CD8-V450 (2.5μl, BD 561426). and incubated on ice for 15 minutes. After incubation, 150μl FACS Wash was added. Cells were pelleted at 1500rpm for 5 minutes followed by two washes with FACS Wash. Cells were collected in 200μl FACS Wash for flow cytometry analysis. Cells were analyzed using an LSR Fortessa X-20 cell analyzer (BD Biosciences). BD CompBeads (51-90-9001229) were used to calculate compensation for all antibody stains and methanol-fixed Namalwa cells (ATCC CRL1432) +/-KSHV were used to calculate compensation for GFP and the fixable viability stain. Flow cytometry data was analyzed using FlowJo software and exported for quantitative analysis in R as described below.

### ICCS for IL-21 secretion

At 3dpi, cultures were treated for 6 hours with [4 ul for every 6ml of cel culture] monensin to block cytokine secretion. Following incubation, approximately 1 million cells were harvested and viability and surface staining for T cell lineage markers was performed as described above. After the final wash, cells were fixed for 10 minutes in BD cytofix/cytoperm (51-2090KZ), pelleted and further treated for 10 minutes with cytofix/cytoperm+10% DMSO (superperm) to more effectively get intracellular antibodies into the nucleus. Intracellular antibodies, as follows, were diluted in 1x BD Permwash (51-2091KZ) and left on fixed cells overnight at 4°C. RoR-γT-BV421 (563282, 5μl/test), FoxP3-BB700 (566527, 5 μl/test), IL-21-APC (513007, 5 μl/test), BLC6-BV711 (561080, 5μl/test). Cells were then washed twice with 1x permwash and analyzed as described above.

### RT-PCR

At 3 days post infection, 1(10)^6^ lymphocytes were harvested into an equal volume of Trizol and DNA/RNA shield (Zymo Research R110-250). Total RNA was extracted using using Zymo Directzol Microprep (Zymo Research R2060) according to manufacturer instructions. RNA was eluted in 10μl H2O containing 2U RNase inhibitors and a second DNase step was performed for 30 minutes using the Turbo DNA-Free kit (Invitrogen AM1907M) according to manufacturer instructions. One-step RT-PCR cDNA synthesis and preamplification of GAPDH, LANA and K8.1 transcripts was performed on 15ng of total RNA using the Superscript III One-step RT-PCR kit (ThermoFisher 12574026).

Duplicate no RT (NRT) control reactions were assembled for each sample containing only Platinum Taq DNA polymerase (Thermofisher 15966005) instead of the Superscript III RT/Taq DNA polymerase mix. After cDNA synthesis and 20 cycles of target pre-amplification, 2μl of pre-amplified cDNA or NRT control reaction was used as template for multiplexed real-time PCR reactions using TaqProbe 5x qPCR MasterMix-Multiplex (ABM MasterMix-5PM), 5% DMSO, primers at 900nM and probes at 250nM against target genes. All primer and probe sequences used in these assays have been previously published [10]. Real time PCR was performed using a 40-cycle program on a Biorad real time thermocycler. Data is represented as quantitation cycle (Cq) and assays in which there was no detectable Cq value were set numerically as Cq = 41 for analysis and data visualization. The expression of each gene was normalized to that of a housekeeping gene *GAPDH*.

### Statistical Analysis

The indicated data sets and statistical analysis were performed in Rstudio software using ggplot2 [41], ggcorrplot [42], ggally [43] and tidyverse [44] packages. Statistical analysis was performed using rstatix [45] package. Specific methods of statistical analysis including Anova, independent t-test and Pearson correlations and resulting values for significance and correlation are detailed in the corresponding figure legends.

## Supplemental Materials

**Supplemental Table 1A**: One-way repeated measures ANOVA for the effect of IL-21 treatment on GFP distribution in B cell subsets. Sorted by p-value.

**Supplemental Table 1B**: Two-way repeated measures ANOVA for the effect of KSHV infection (Cond) and IL-21 treatment (Tx) on total GFP and frequencies of B cell subsets. Sorted by p-value.

**Supplemental Table 2A**: One-way repeated measures ANOVA analysis for the dose effect of IL-21 neutralizing antibody on GFP frequency within B cell subsets. Sorted by p value.

**Supplemental Table 2B**: One-way repeated measures ANOVA analysis for the dose effect of IL-21 neutralizing antibody on B cell subset frequencies. Sorted by p value.

**Supplemental Table 3A**: Pairwise correlations using Pearson’s method between total GFP+ cells in KSHV-infected conditions at 3 dpi and the baseline (0 dpi) frequency of each B cell subset within IL-21R+ cells. Sorted by p value.

**Supplemental Table 3B**: Pairwise correlations using Pearson’s method between the change in GFP+ cells in response to 100ng/ml IL-21 treatment (GFP+ Treatment-GFP+ Control) in KSHV-infected conditions at 3 dpi and the baseline (0 dpi) frequency of each B cell subset within IL-21R+ cells. Sorted by p value.

**Supplemental Table 3C**: Pairwise correlations using Pearson’s method between the change in frequency of plasma cells with IL-21 treatment at 3dpi (PC treated – PC control) in KSHV-infected conditions at 3 dpi and the baseline (0 dpi) frequency of each B cell subset within IL-21R+ cells. Sorted by p value.

**Supplemental Table 4A**: Two-way repeated measures ANOVA analysis for the effect of IL-21 treatment (Tx) and Baseline vs. Mock vs. KSHV infection (Cond) on the frequency of B cell subsets within IL-21R+ at 3 dpi. Sorted by p value.

**Supplemental Table 4B**: Post-hoc paired T-test for the Cond effects in the ANOVA analysis shown in Table 4A. Significance indicates differences between Baseline (BL), Mock and KSHV-infected cultures grouped by IL-21 treatment (Tx); NT=untreated, IL21=100ng/ml IL-21 treated. Sorted by adjusted p-value using Holm correction for multiple comparisons.

**Supplemental Table 4C**: Post-hoc paired T-test for the Tx effects in the ANOVA analysis shown in Table 4A. Significance indicates differences between NT=untreated, IL21=100ng/ml IL-21 treated grouped by infection condition (Mock or KSHV-infected). Sorted by p-value.

**Supplemental Table 4D**: Two-way repeated measures ANOVA analysis for the effect of IL-21 treatment (TX) and Baseline vs. Mock vs. KSHV infection (Cond) on the MFI of IL-21R within IL-21R+ B cell subsets at 3 dpi. Sorted by p value.

**Supplemental Table 5A**: Pairwise correlations using Pearson’s method between T cell frequencies within IL-21+ T cells and total GFP+ B lymphocytes at 3 dpi. Sorted by p value.

**Supplemental Table 5B**: Pairwise correlations using Pearson’s method between the iMFI of IL-21 for T cell subsets and total GFP+ B lymphocytes at 3 dpi. Sorted by p value.

**Supplemental Table 5C**: Pairwise correlations using Pearson’s method between the baseline (Day 0) frequencies T cell subsets and total GFP+ B lymphocytes at 3 dpi. Sorted by p value.

**Supplemental Figure 1**: Representative gating scheme for (A) B cell and (B) T cell immunophenotyping using lineage definitions as detailed in Table 2 (C) Full panel and FMO control for ICCS staining of IL-21 in the T cell immunophenotyping panel.

**Supplemental Figure 2**: (A) Total T cell subset frequencies in Mock and KSHV-infected cultures as in Figure 6. Correlograms of Pearson correlations between the distribution of KSHV infection within B cell subsets (y-axis) and (B) total IL-21 or the contribution of individual T cell subsets to IL-21 secretion at 3 dpi (x-axis), (C) baseline T cell frequencies, and (D) iMFI of IL-21 within T cell subsets at 3dpi. Pearson’s r values with an absolute value greater than or equal to 0.53 are statistically significant for this dataset.

## References

1. Chang, Y., et al., Identification of herpesvirus-like DNA sequences in AIDS-associated Kaposi’s sarcoma. Science, 1994. 266(5192): p. 1865–1869.

2. Ablashi, D.V., et al., Spectrum of Kaposi’s sarcoma-associated herpesvirus, or human herpesvirus 8, diseases. Clinical microbiology reviews, 2002. 15(3): p. 439–464.

3. Cesarman, E., et al., Kaposi’s sarcoma–associated herpesvirus-like DNA sequences in AIDS-related body-cavity–based lymphomas. New England Journal of Medicine, 1995. 332(18): p. 1186–1191.

4. Soulier, J., et al., Kaposi’s sarcoma-associated herpesvirus-like DNA sequences in multicentric Castleman’s disease [see comments]. Blood, 1995. 86(4): p. 1276–1280.

5. Uldrick, T.S., et al., An interleukin-6-related systemic inflammatory syndrome in patients co-infected with Kaposi sarcoma-associated herpesvirus and HIV but without Multicentric Castleman disease. Clinical Infectious Diseases, 2010. 51(3): p. 350–358.

6. Bouvard, V., et al., A review of human carcinogens--Part B: biological agents. The Lancet. Oncology, 2009. 10(4): p. 321–322.

7. Parkin, D.M., The global health burden of infection-associated cancers in the year 2002. International journal of cancer, 2006. 118(12): p. 3030–3044.

8. Ganem, D., KSHV and the pathogenesis of Kaposi sarcoma: listening to human biology and medicine. The Journal of clinical investigation, 2010. 120(4): p. 939–949.

9. Casper, C., et al., Frequent and asymptomatic oropharyngeal shedding of human herpesvirus 8 among immunocompetent men. The Journal of infectious diseases, 2007. 195(1): p. 30–36.

10. Aalam, F., et al., Analysis of KSHV B lymphocyte lineage tropism in human tonsil reveals efficient infection of CD138+ plasma cells. PLoS pathogens, 2020. 16(10): p. e1008968.

11. Alomari, N. and J. Totonchy, Cytokine-Targeted Therapeutics for KSHV-Associated Disease. Viruses, 2020. 12(10): p. 1097.

12. Parrish-Novak, J., et al., Interleukin-21 and the IL-21 receptor: novel effectors of NK and T cell responses. Journal of leukocyte biology, 2002. 72(5): p. 856–863.

13. Parrish-Novak, J., et al., Interleukin 21 and its receptor are involved in NK cell expansion and regulation of lymphocyte function. Nature, 2000. 408(6808): p. 57–63.

14. Kuchen, S., et al., Essential role of IL-21 in B cell activation, expansion, and plasma cell generation during CD4+ T cell-B cell collaboration. The journal of immunology, 2007. 179(9): p. 5886–5896.

15. Konforte, D., N. Simard, and C.J. Paige, IL-21: an executor of B cell fate. The Journal of Immunology, 2009. 182(4): p. 1781–1787.

16. Ozaki, K., et al., Regulation of B cell differentiation and plasma cell generation by IL-21, a novel inducer of Blimp-1 and Bcl-6. The Journal of Immunology, 2004. 173(9): p. 5361–5371.

17. Calame, K.L., K.-I. Lin, and C. Tunyaplin, Regulatory mechanisms that determine the development and function of plasma cells. Annual review of immunology, 2003. 21(1): p. 205–230.

18. Schmitz, I., et al., IL-21 restricts virus-driven Treg cell expansion in chronic LCMV infection. PLoS pathogens, 2013. 9(5): p. e1003362.

19. Pallikkuth, S., et al., Upregulation of IL-21 receptor on B cells and IL-21 secretion distinguishes novel 2009 H1N1 vaccine responders from nonresponders among HIV-infected persons on combination antiretroviral therapy. The Journal of Immunology, 2011. 186(11): p. 6173–6181.

20. Collins, C.M. and S.H. Speck, Interleukin 21 signaling in B cells is required for efficient establishment of murine gammaherpesvirus latency. PLoS pathogens, 2015. 11(4): p. e1004831.

21. Konforte, D. and C.J. Paige, Identification of cellular intermediates and molecular pathways induced by IL-21 in human B cells. The Journal of Immunology, 2006. 177(12): p. 8381–8392.

22. Konforte, D., N. Simard, and C.J. Paige, Interleukin-21 regulates expression of key Epstein–Barr virus oncoproteins, EBNA2 and LMP1, in infected human B cells. Virology, 2008. 374(1): p. 100–113.

23. Yajima, H., et al., Loss of interleukin-21 leads to atrophic germinal centers in multicentric Castleman’s disease. Annals of hematology, 2016. 95(1): p. 35–40.

24. Gasperini, P. and G. Tosato, Targeting the mammalian target of Rapamycin to inhibit VEGF and cytokines for the treatment of primary effusion lymphoma. Leukemia, 2009. 23(10): p. 1867–1874.

25. Gasperini, P., S. Sakakibara, and G. Tosato, Contribution of viral and cellular cytokines to Kaposi’s sarcoma-associated herpesvirus pathogenesis. Journal of leukocyte biology, 2008. 84(4): p. 994–1000.

26. Totonchy, J., et al., KSHV induces immunoglobulin rearrangements in mature B lymphocytes. PLoS pathogens, 2018. 14(4): p. e1006967.

27. Darrah, P.A., et al., Multifunctional TH1 cells define a correlate of vaccine-mediated protection against Leishmania major. Nature medicine, 2007. 13(7): p. 843–850.

28. Chang, J., et al., Induction of Kaposi’s sarcoma-associated herpesvirus from latency by inflammatory cytokines. Virology, 2000. 266: p. 17–25.

29. Chang, J., et al., Inflammatory cytokines and the reactivation of Kaposi’s sarcoma-associated herpesvirus lytic replication. Virology, 2000. 266(1): p. 17–25.

30. Gantt, S., et al., Prospective characterization of the risk factors for transmission and symptoms of primary human herpesvirus infections among Ugandan infants. The Journal of infectious diseases, 2016. 214(1): p. 36–44.

31. Jondle, C., et al., Gammaherpesvirus Usurps Host IL-17 Signaling To Support the Establishment of Chronic Infection. Mbio, 2021. 12(2): p. e00566–21.

32. Jondle, C., et al., T Cell-Intrinsic Interferon Regulatory Factor 1 Expression Suppresses Differentiation of CD4+ T Cell Populations That Support Chronic Gammaherpesvirus Infection. Journal of Virology, 2021. 95(20): p. e00726–21.

33. Gaddi, P.J. and G.S. Yap, Cytokine regulation of immunopathology in toxoplasmosis. Immunology and cell biology, 2007. 85(2): p. 155–159.

34. Kelly, M.N., et al., Interleukin-17/interleukin-17 receptor-mediated signaling is important for generation of an optimal polymorphonuclear response against Toxoplasma gondii infection. Infection and immunity, 2005. 73(1): p. 617–621.

35. Shainheit, M.G., et al., The pathogenic Th17 cell response to major schistosome egg antigen is sequentially dependent on IL-23 and IL-1β. The Journal of Immunology, 2011. 187(10): p. 5328–5335.

36. Wen, X., et al., Dynamics of Th17 cells and their role in Schistosoma japonicum infection in C57BL/6 mice. PLoS neglected tropical diseases, 2011. 5(11): p. e1399.

37. Wakeham, K., et al., Parasite infection is associated with Kaposi’s sarcoma associated herpesvirus (KSHV) in Ugandan women. Infectious agents and cancer, 2011. 6(1): p. 1–7.

38. Totonchy, J., Extrafollicular activities: perspectives on HIV infection, germinal center-independent maturation pathways, and KSHV-mediated lymphoproliferation. Current opinion in virology, 2017. 26: p. 69–73.

39. Quigley, M.F., et al., CXCR5+ CCR7–CD8 T cells are early effector memory cells that infiltrate tonsil B cell follicles. European journal of immunology, 2007. 37(12): p. 3352–3362.

40. Yang, R., et al., IL-6 promotes the differentiation of a subset of naive CD8+ T cells into IL-21–producing B helper CD8+ T cells. Journal of Experimental Medicine, 2016. 213(11): p. 2281–2291.

41. Wickham, H., Ggplot2: elegant graphics for data analysis. Use R! 2009, New York: Springer. viii, 212 p.

42. Kassambara, A., ggcorrplot: Visualization of a Correlation Matrix using ‘ggplot2′ 2019.

43. Schloerke, B., J. Crowley, and D. Cook, Package ‘GGally’. Extension to ‘ggplot2.’See, 2018. 713.

44. Lander, J.P., R for everyone: advanced analytics and graphics. Second edition ed. The Addison Wesley data and anlytics series. 2017, Boston: Addison-Wesley. xxiv, 528 pages.

45. Kassambara, A., rstatix: Pipe-Friendly Framework for Basic Statistical Tests. 2020.

