## Supplemental Tables and Figures for "IL-21 signaling promotes the establishment of KSHV infection in human tonsil lymphocytes by increasing early targeting of plasma cells"

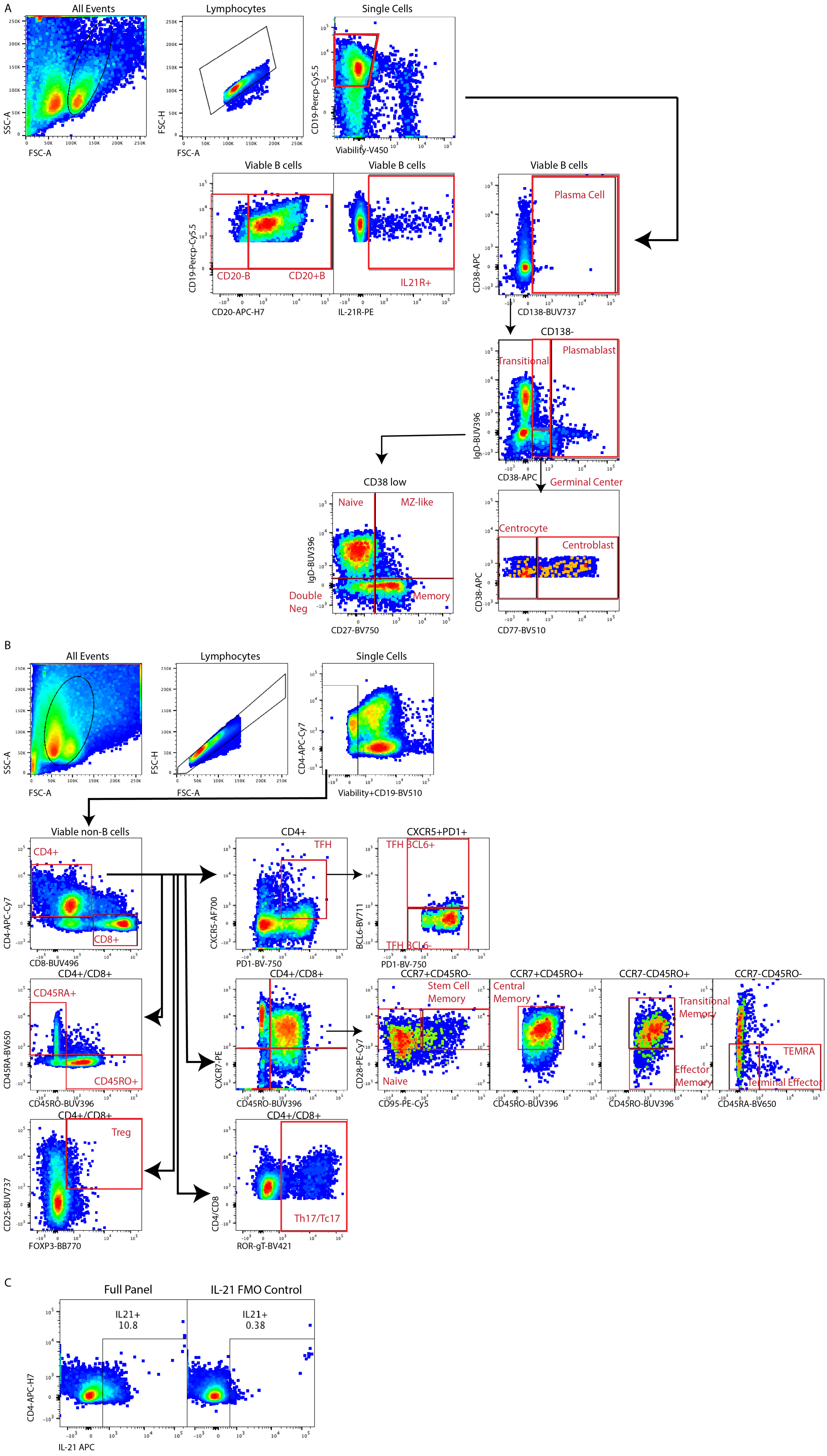
A
All Events
Lymphocytes
Single Cells
SSC-A
FSC-A
FSC-H
CD19-Percp-Cy5.5
Viability-V450
Viable B cells
CD20-B
CD20+B
CD20-APC-H7
IL21R+
IL-21R-PE
Plasma Cell
CD38-APC
CD138-BUV737
CD138-
Transitional
Plasmablast
IgD-BUV396
Germinal Center
CD38 low
Naive
MZ-like
Double Neg
Memory
CD27-BV750
Centrocyte
Centroblast
CD77-BV510
B
Viability+CD19-BV510
CD4-APC-Cy7
Viable non-B cells
CD4+
CD8+
CD8-BUV496
CXCR5-AF700
TFH
PD1-BV-750
CXCR5+PD1+
TFH BCL6+
TFH BCL6-
BCL6-BV711
CD4+/CD8+
CD45RA+
CD45RO+
CD45RA-BV650
CD45RO-BUV396
CXCR7-PE
CCR7+CD45RO-
Stem Cell Memory
Naive
CD28-PE-Cy7
CD95-PE-Cy5
CCR7+CD45RO+
Central Memory
CCR7-CD45RO+
Transitional Memory
Effector Memory
CCR7-CD45RO-
TEMRA
Terminal Effector
Treg
CD25-BUV737
FOXP3-BB770
Th17/Tc17
CD4/CD8
ROR-gT-BV421
C
Full Panel
IL-21 FMO Control
IL21+
10.8
IL21+
0.38
CD4-APC-H7
IL-21 APC

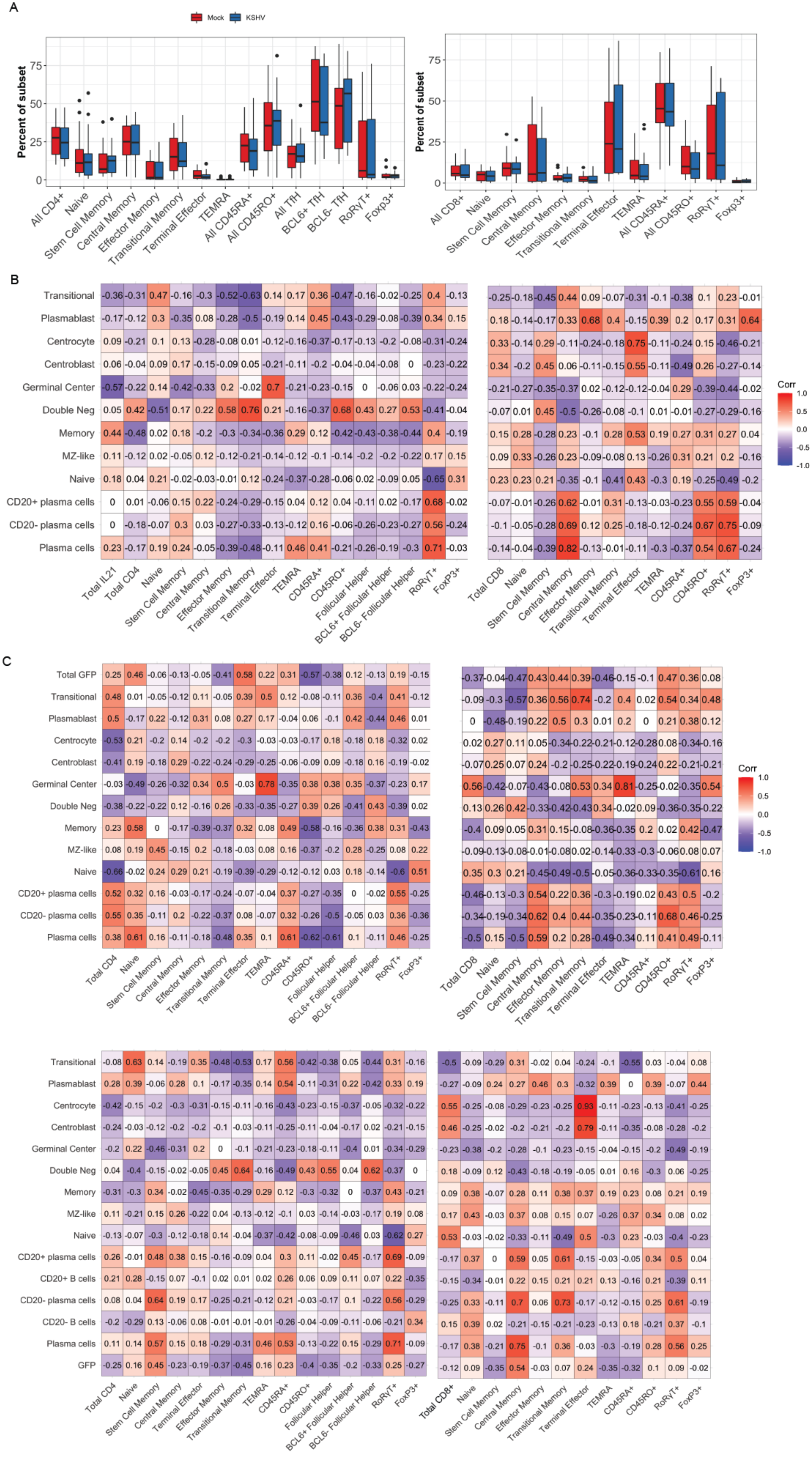
A
Mock
KSHV
Percent of subset
All CD4+
Naive
Stem Cell Memory
Central Memory
Effector Memory
Transitional Memory
Terminal Effector
TEMRA
All CD45RA+
All CD45RO+
All TfH
BCL6+ TfH
BCL6- TfH
RoRγT+
Foxp3+
All CD8+
B
Transitional
Plasmablast
Centrocyte
Centroblast
Germinal Center
Double Neg
Memory
MZ-like
Naive
CD20+ plasma cells
CD20- plasma cells
Plasma cells
Total IL21
Total CD4
CD45RA+
CD45RO+
Follicular Helper
BCL6+ Follicular Helper
BCL6- Follicular Helper
RoRγT+
FoxP3+
Total CD8
Corr
C
Total GFP
CD20+ B cells
CD20- B cells
GFP
Total CD8+

**Supplemental Table 1A**: One-way repeated measures ANOVA for the effect of IL-21 treatment on GFP distribution in B cell subsets. Sorted by p-value.

| **Percent GFP+ within subset** | **Effect** | **DFn** | **DFd** | **F** | **p** | **p<.05** | **ges** |
|---|---|---|---|---|---|---|---|
| Plasma cell | Tx | 1 | 13 | 6.633 | 0.023 | * | 0.133 |
| Plasmablast | Tx | 1 | 13 | 4.333 | 0.058 | | 0.049 |
| Germinal center | Tx | 1 | 13 | 3.973 | 0.068 | | 0.022 |
| MZ-like | Tx | 1 | 13 | 2.556 | 0.134 | | 0.026 |
| Naive | Tx | 1 | 13 | 2.036 | 0.177 | | 0.043 |
| Transitional | Tx | 1 | 13 | 1.121 | 0.309 | | 0.008 |
| Memory | Tx | 1 | 13 | 0.061 | 0.808 | | 0.001 |
| Double Negative | Tx | 1 | 13 | 0.039 | 0.847 | | 3e-04 |

**Supplemental Table 1B**: Two-way repeated measures ANOVA for the effect of KSHV infection (Cond) and IL-21 treatment (Tx) on total GFP and frequencies of B cell subsets. Sorted by p-value.

| **Frequency within Viable B** | **Effect** | **DFn** | **DFd** | **F** | **p** | **p<.05** | **ges** |
|---|---|---|---|---|---|---|---|
| GFP | Cond | 1 | 13 | 102.896 | 1.52e-07 | * | 0.708 |
| Plasma Cell | Tx:Cond | 1 | 13 | 22.779 | 0.000364 | * | 0.09 |
| Plasma Cell | Tx | 1 | 13 | 18.596 | 0.000844 | * | 0.163 |
| Plasma Cell | Cond | 1 | 13 | 18.575 | 0.000848 | * | 0.166 |
| Naive | Tx | 1 | 13 | 13.643 | 0.003 | * | 0.099 |
| CD20- Plasma Cell | Tx:Cond | 1 | 13 | 12.645 | 0.004 | * | 0.043 |
| Naive | Cond | 1 | 13 | 8.462 | 0.012 | * | 0.069 |
| CD20- Plasma Cell | Tx:Cond | 1 | 13 | 7.798 | 0.015 | * | 0.012 |
| GFP | Tx:Cond | 1 | 13 | 7.635 | 0.016 | * | 0.084 |
| GFP | Tx | 1 | 13 | 6.425 | 0.025 | * | 0.081 |
| CD20+ B cell | Cond | 1 | 13 | 6.234 | 0.027 | * | 0.016 |
| Plasmablast | Cond | 1 | 13 | 6.023 | 0.029 | * | 0.02 |
| CD20- Plasma Cell | Tx | 1 | 13 | 4.858 | 0.046 | * | 0.028 |
| CD20+ B cell | Tx | 1 | 13 | 4.563 | 0.052 | | 0.032 |
| Plasmablast | Tx | 1 | 13 | 4.115 | 0.064 | | 0.016 |
| CD20- B cell | Cond | 1 | 13 | 3.949 | 0.068 | | 0.011 |
| MZ-like | Tx | 1 | 13 | 3.897 | 0.07 | | 0.023 |
| CD20- Plasma Cell | Cond | 1 | 13 | 3.875 | 0.071 | | 0.018 |
| CD20- B cell | Tx | 1 | 13 | 3.663 | 0.078 | | 0.03 |
| Transitional | Cond | 1 | 13 | 3.339 | 0.091 | | 0.011 |
| CD20+ B cell | Tx:Cond | 1 | 13 | 3.106 | 0.101 | | 0.008 |
| MZ-like | Tx:Cond | 1 | 13 | 3.08 | 0.103 | | 0.019 |
| Memory | Cond | 1 | 13 | 2.764 | 0.12 | | 0.046 |
| Germinal Center | Tx | 1 | 13 | 2.472 | 0.14 | | 0.007 |
| Plasmablast | Tx:Cond | 1 | 13 | 2.113 | 0.17 | | 0.003 |
| CD20- B cell | Tx:Cond | 1 | 13 | 1.66 | 0.22 | | 0.006 |
| all CD19+ | Tx:Cond | 1 | 13 | 1.246 | 0.284 | | 0.003 |

| | | | | | | | |
|---|---|---|---|---|---|---|---|
| Double Neg | Tx:Cond | 1 | 13 | 1.082 | 0.317 | | 0.004 |
| Naive | Tx:Cond | 1 | 13 | 0.966 | 0.344 | | 0.002 |
| Transitional | Tx:Cond | 1 | 13 | 0.857 | 0.371 | | 0.002 |
| Germinal Center | Cond | 1 | 13 | 0.74 | 0.405 | | 0.004 |
| Double Neg | Tx | 1 | 13 | 0.734 | 0.407 | | 0.004 |
| all CD19+ | Cond | 1 | 13 | 0.695 | 0.42 | | 0.004 |
| Transitional | Tx | 1 | 13 | 0.475 | 0.503 | | 0.000824 |
| all CD19+ | Tx | 1 | 13 | 0.375 | 0.551 | | 0.001 |
| CD20- Plasma Cell | Tx | 1 | 13 | 0.366 | 0.555 | | 0.004 |
| Germinal Center | Tx:Cond | 1 | 13 | 0.328 | 0.577 | | 0.001 |
| MZ-like | Cond | 1 | 13 | 0.283 | 0.604 | | 0.000917 |
| Memory | Tx | 1 | 13 | 0.274 | 0.61 | | 0.001 |
| Double Neg | Cond | 1 | 13 | 0.202 | 0.661 | | 0.002 |
| Memory | Tx:Cond | 1 | 13 | 0.111 | 0.744 | | 0.000299 |
| CD20- Plasma Cell | Cond | 1 | 13 | 0.011 | 0.916 | | 0.000149 |

**Supplemental Table 2A**: One-way repeated measures ANOVA analysis for the dose  effect of IL-21 neutralizing antibody on GFP frequency within B cell subsets. Sorted by p value.

| **Percent GFP+ within subset** | Effect | DFn | DFd | F | p | p<.05 | ges |
|---|---|---|---|---|---|---|---|
| Transitional | Dose | 3 | 30 | 4.036 | 0.016 | * | 0.043 |
| All Plasma Cells | Dose | 3 | 30 | 2.202 | 0.108 | | 0.05 |
| Plasmablast | Dose | 3 | 30 | 2.026 | 0.131 | | 0.025 |
| Memory | Dose | 3 | 30 | 1.999 | 0.135 | | 0.041 |
| centroblast | Dose | 3 | 30 | 1.754 | 0.177 | | 0.077 |
| Naive | Dose | 3 | 30 | 1.416 | 0.257 | | 0.018 |
| Germinal Center | Dose | 3 | 30 | 1.411 | 0.259 | | 0.018 |
| MZ-like | Dose | 1.62 | 16.2 | 1.423 | 0.266 | | 0.027 |
| Double Neg | Dose | 3 | 30 | 1.209 | 0.323 | | 0.019 |
| centrocyte | Dose | 3 | 30 | 0.861 | 0.472 | | 0.012 |

**Supplemental Table 2B**: One-way repeated measures ANOVA analysis for the dose effect of IL-21 neutralizing antibody on B cell subset frequencies. Sorted by p value.

| **Percent subset within Viable B** | Effect | DFn | DFd | F | p | p<.05 | ges |
|---|---|---|---|---|---|---|---|
| GFP | Dose | 3 | 30 | 9.426 | 0.000152 | * | 0.31 |
| All Plasma Cells | Dose | 1.64 | 16.38 | 2.146 | 0.154 | | 0.064 |
| CD20+ B cell | Dose | 1.1 | 10.99 | 1.883 | 0.199 | | 0.01 |
| Germinal Center | Dose | 1.78 | 17.81 | 1.435 | 0.263 | | 0.009 |
| centrocyte | Dose | 1.68 | 16.8 | 1.377 | 0.276 | | 0.009 |
| CD20- B cell | Dose | 1.07 | 10.69 | 1.313 | 0.281 | | 0.015 |
| Plasmablast | Dose | 3 | 30 | 1.196 | 0.328 | | 0.008 |
| Transitional | Dose | 3 | 30 | 1.197 | 0.328 | | 0.011 |
| MZ-like | Dose | 1.66 | 16.56 | 1.082 | 0.35 | | 0.009 |
| CD20- Plasma Cell | Dose | 3 | 30 | 1.048 | 0.386 | | 0.026 |
| Double Neg | Dose | 3 | 30 | 0.905 | 0.45 | | 0.021 |
| CD20- Plasma Cell | Dose | 3 | 30 | 0.839 | 0.483 | | 0.024 |
| Naive | Dose | 2.03 | 20.25 | 0.534 | 0.597 | | 0.008 |

| | | | | | | | |
|---|---|---|---|---|---|---|---|
| Memory | Dose | 3 | 30 | 0.592 | 0.625 | | 0.015 |
| centroblast | Dose | 1.19 | 11.92 | 0.227 | 0.685 | | 0.006 |
| all CD19+ | Dose | 1.62 | 16.2 | 0.259 | 0.729 | | 0.005 |

**Supplemental Table 3A**: Pairwise correlations using Pearson's method between total GFP+ cells in KSHV-infected conditions at 3 dpi and the baseline (0 dpi) frequency of each B cell subset within IL-21R+ cells. Sorted by p value.

| **Frequency of Subset within IL-21R+ at Day 0** | **Cor to total GFP** | statistic | p | p>.05 | conf.low | conf.high |
|---|---|---|---|---|---|---|
| Plasmablast | 0.81 | 4.65 | 0.000697 | *** | 0.48 | 0.94 |
| Naive | -0.57 | -2.29 | 0.0424 | * | -0.85 | -0.03 |
| Transitional | 0.39 | 1.40 | 0.19 | | -0.21 | 0.77 |
| Memory | -0.38 | -1.37 | 0.198 | | -0.77 | 0.21 |
| CD20- Plasma Cell | 0.38 | 1.36 | 0.2 | | -0.22 | 0.77 |
| Germinal Center | -0.38 | -1.35 | 0.203 | | -0.77 | 0.22 |
| CD20+ B cell | -0.36 | -1.30 | 0.221 | | -0.76 | 0.23 |
| CD20- B cells | 0.34 | 1.201 | 0.255 | | -0.26 | 0.75 |
| Plasma cell | 0.25 | 0.87 | 0.402 | | -0.35 | 0.71 |
| CD20+ Plasma Cell | 0.23 | 0.80 | 0.441 | | -0.36 | 0.70 |
| Total IL21R+ B cells | 0.21 | 0.71 | 0.494 | | -0.39 | 0.68 |
| Double Neg | -0.18 | -0.60 | 0.56 | | -0.66 | 0.41 |
| MZ-like | -0.075 | -0.25 | 0.808 | | -0.61 | 0.50 |

**Supplemental Table 3B**: Pairwise correlations using Pearson's method between the change in GFP+ cells in response to 100ng/ml IL-21 treatment (GFP+ Treatment- GFP+ Control) in KSHV-infected conditions at 3 dpi and the baseline (0 dpi) frequency of each B cell subset within IL-21R+ cells. Sorted by p value.

| **Frequency of Subset within IL-21R+ at Day 0** | **Cor to ΔGFP** | statistic | p | p>.05 | conf.low | conf.high |
|---|---|---|---|---|---|---|
| Naive | 0.82 | 3.45 | 0.0137 | ** | 0.30 | 0.97 |
| Memory | 0.71 | 2.48 | 0.0477 | * | 0.01 | 0.94 |
| CD20+ Plasma Cells | -0.59 | -1.81 | 0.12 | | -0.92 | 0.19 |
| PC | -0.57 | -1.69 | 0.143 | | -0.91 | 0.23 |
| Plasmablast | -0.57 | -1.68 | 0.143 | | -0.91 | 0.23 |
| Germinal Center | 0.53 | 1.51 | 0.181 | | -0.28 | 0.90 |
| Transitional | -0.46 | -1.27 | 0.251 | | -0.88 | 0.36 |
| Double Neg | -0.46 | -1.26 | 0.254 | | -0.88 | 0.36 |
| MZ-like | 0.43 | 1.18 | 0.283 | | -0.39 | 0.87 |
| CD20- Plasma Cell | -0.36 | -0.95 | 0.38 | | -0.85 | 0.46 |
| CD20+ B cell | 0.19 | 0.48 | 0.649 | | -0.59 | 0.79 |
| CD20- B cells | -0.13 | -0.33 | 0.75 | | -0.77 | 0.63 |
| Total IL21R+ B cells | -0.13 | -0.31 | 0.766 | | -0.76 | 0.64 |

**Supplemental Table 3C**: Pairwise correlations using Pearson's method between the change in frequency of plasma cells with IL-21 treatment at 3dpi (PC treated – PC control) in KSHV-infected conditions at 3 dpi and the baseline (0 dpi) frequency of each B cell subset within IL-21R+ cells. Sorted by p value.

| **Frequency of Subset within IL-21R+ at Day 0** | **Cor to ΔPC** | statistic | p | P<.05 | conf.low | conf.high |
|---|---|---|---|---|---|---|
| Double Neg | -0.71 | -2.50 | 0.0464 | * | -0.94 | -0.02 |
| CD20+ Plasma Cells | -0.6 | -1.83 | 0.117 | | -0.92 | 0.18 |
| PC | -0.56 | -1.68 | 0.145 | | -0.91 | 0.23 |
| MZ-like | 0.17 | 0.43 | 0.686 | | -0.61 | 0.78 |
| Germinal Center | 0.16 | 0.40 | 0.701 | | -0.61 | 0.78 |
| Total IL21R+ B cells | -0.13 | -0.31 | 0.766 | | -0.76 | 0.63 |
| Plasmablast | 0.13 | 0.31 | 0.766 | | -0.644 | 0.76 |
| CD20+ B cell | 0.11 | 0.27 | 0.795 | | -0.64 | 0.761 |
| CD20- Plasma Cell | -0.098 | -0.24 | 0.817 | | -0.75 | 0.65 |
| CD20- B cells | -0.074 | -0.18 | 0.862 | | -0.74 | 0.67 |
| Transitional | 0.071 | 0.18 | 0.867 | | -0.67 | 0.74 |
| Naive | -0.056 | -0.14 | 0.895 | | -0.73 | 0.68 |
| Memory | 0.031 | 0.08 | 0.942 | | -0.69 | 0.72 |

**Supplemental Table 4A**: Two-way repeated measures ANOVA analysis for the effect of IL-21 treatment (Tx) and Baseline vs. Mock vs. KSHV infection (Cond) on the frequency of B cell subsets within IL-21R+ at 3 dpi. Sorted by p value.

| **Percent of subset within IL-21R+ at 3dpi** | Effect | DFn | DFd | F | p | p<.05 | ges |
|---|---|---|---|---|---|---|---|
| Memory | Tx | 1 | 9 | 15.641 | 0.003 | * | 0.017 |
| Plasmablast | Cond | 1 | 9 | 8.826 | 0.016 | * | 0.079 |
| Plasmablast | Tx | 1 | 9 | 4.974 | 0.053 | | 0.09 |
| Plasma cell | Tx | 1 | 9 | 3.559 | 0.092 | | 0.049 |
| MZ-like | Tx | 1 | 9 | 2.987 | 0.118 | | 0.06 |
| MZ-like | Cond | 1 | 9 | 2.706 | 0.134 | | 0.042 |
| Plasma cell | Cond | 1 | 9 | 2.521 | 0.147 | | 0.012 |
| Naive | Tx:Cond | 1 | 9 | 2.173 | 0.175 | | 0.004 |
| Double Neg | Tx:Cond | 1 | 9 | 2.054 | 0.186 | | 0.017 |
| Memory | Cond | 1 | 9 | 1.177 | 0.306 | | 0.008 |
| Memory | Tx:Cond | 1 | 9 | 1.089 | 0.324 | | 0.009 |
| Double Neg | Tx | 1 | 9 | 1.079 | 0.326 | | 0.005 |
| Germinal Center | Tx | 1 | 9 | 0.807 | 0.392 | | 0.005 |
| Germinal Center | Cond | 1 | 9 | 0.576 | 0.467 | | 0.006 |
| MZ-like | Tx:Cond | 1 | 9 | 0.503 | 0.496 | | 0.005 |
| Transitional | Cond | 1 | 9 | 0.477 | 0.507 | | 0.004 |
| Naive | Cond | 1 | 9 | 0.395 | 0.546 | | 0.000966 |
| Transitional | Tx:Cond | 1 | 9 | 0.374 | 0.556 | | 0.002 |
| Plasma cell | Tx:Cond | 1 | 9 | 0.349 | 0.569 | | 0.003 |

| | | | | | | | |
|---|---|---|---|---|---|---|---|
| Plasmablast | Tx:Cond | 1 | 9 | 0.326 | 0.582 | | 0.002 |
| Double Neg | Cond | 1 | 9 | 0.111 | 0.747 | | 0.000281 |
| Germinal Center | Tx:Cond | 1 | 9 | 0.095 | 0.765 | | 0.000843 |
| Memory | Tx | 1 | 9 | 0.054 | 0.821 | | 0.00065 |
| Transitional | Tx | 1 | 9 | 0.01 | 0.922 | | 9.11e-05 |

**Supplemental Table 4B**: Post-hoc paired T-test for the Cond effects in the ANOVA analysis shown in Table 4A. Significance indicates differences between Baseline (BL), Mock and KSHV-infected cultures grouped by IL-21 treatment (Tx); NT=untreated, IL21=100ng/ml IL-21 treated. Sorted by adjusted p-value using Holm correction for multiple comparisons.

| Tx | **Percent of subset within IL-21R+ at 3dpi** | group1 | group2 | n1 | n2 | statistic | df | p | p.adj | p.adj. signif |
|---|---|---|---|---|---|---|---|---|---|---|
| NT | Transitional | BL | KSHV | 10 | 10 | -4.54 | 9 | 0.001 | 0.003 | ** |
| NT | Transitional | BL | Mock | 10 | 10 | -4.72 | 9 | 0.001 | 0.003 | ** |
| NT | Plasmablast | BL | Mock | 10 | 10 | 3.17 | 9 | 0.011 | 0.034 | * |
| NT | Plasmablast | KSHV | Mock | 10 | 10 | 3.04 | 9 | 0.014 | 0.034 | * |
| NT | MZ-like | BL | KSHV | 10 | 10 | 2.59 | 9 | 0.029 | 0.087 | ns |
| NT | MZ-like | BL | Mock | 10 | 10 | 2.32 | 9 | 0.046 | 0.091 | ns |
| NT | Naive | BL | Mock | 10 | 10 | -2.23 | 9 | 0.053 | 0.159 | ns |
| NT | Plasma Cell | BL | Mock | 10 | 10 | 2.14 | 9 | 0.061 | 0.184 | ns |
| NT | Plasmablast | BL | KSHV | 10 | 10 | 2.12 | 9 | 0.063 | 0.063 | ns |
| IL21 | Plasmablast | KSHV | Mock | 10 | 10 | 2.12 | 9 | 0.064 | 0.064 | ns |
| NT | MZ-like | KSHV | Mock | 10 | 10 | -2.00 | 9 | 0.076 | 0.091 | ns |
| IL21 | Double Neg | KSHV | Mock | 10 | 10 | -1.97 | 9 | 0.081 | 0.081 | ns |
| NT | Naive | BL | KSHV | 10 | 10 | -1.67 | 9 | 0.13 | 0.26 | ns |
| IL21 | Naive | KSHV | Mock | 10 | 10 | 1.65 | 9 | 0.133 | 0.133 | ns |
| NT | Memory | KSHV | Mock | 10 | 10 | -1.42 | 9 | 0.19 | 0.57 | ns |
| NT | Plasma Cell | KSHV | Mock | 10 | 10 | 1.18 | 9 | 0.27 | 0.54 | ns |
| IL21 | Germinal Center | KSHV | Mock | 10 | 10 | -1.12 | 9 | 0.291 | 0.291 | ns |
| NT | Plasma Cell | BL | KSHV | 10 | 10 | 1.08 | 9 | 0.307 | 0.54 | ns |
| NT | Double Neg | BL | KSHV | 10 | 10 | -1.03 | 9 | 0.329 | 0.987 | ns |
| IL21 | Transitional | KSHV | Mock | 10 | 10 | -0.92 | 9 | 0.384 | 0.384 | ns |
| NT | Double Neg | KSHV | Mock | 10 | 10 | 0.90 | 9 | 0.392 | 0.987 | ns |
| IL21 | MZ-like | KSHV | Mock | 10 | 10 | -0.78 | 9 | 0.454 | 0.454 | ns |
| IL21 | Plasma Cell | KSHV | Mock | 10 | 10 | 0.78 | 9 | 0.455 | 0.455 | ns |
| NT | Germinal Center | BL | KSHV | 10 | 10 | 0.64 | 9 | 0.539 | 1 | ns |
| NT | Double Neg | BL | Mock | 10 | 10 | -0.62 | 9 | 0.551 | 0.987 | ns |
| NT | Germinal Center | BL | Mock | 10 | 10 | 0.59 | 9 | 0.569 | 1 | ns |
| NT | Memory | BL | KSHV | 10 | 10 | 0.55 | 9 | 0.595 | 1 | ns |
| NT | Naive | KSHV | Mock | 10 | 10 | -0.48 | 9 | 0.641 | 0.641 | ns |
| NT | Memory | BL | Mock | 10 | 10 | -0.38 | 9 | 0.71 | 1 | ns |
| NT | Germinal Center | KSHV | Mock | 10 | 10 | -0.29 | 9 | 0.777 | 1 | ns |
| NT | Transitional | KSHV | Mock | 10 | 10 | -0.22 | 9 | 0.833 | 0.833 | ns |
| IL21 | Memory | KSHV | Mock | 10 | 10 | 0.03 | 9 | 0.975 | 0.975 | ns |

Supplemental Table 4C: Post-hoc paired T-test for the Tx effects in the ANOVA analysis shown in Table 4A. Significance indicates differences between NT=untreated, IL21=100ng/ml IL-21 treated grouped by infection condition (Mock or KSHV-infected). Sorted by p-value.

| Cond | Percent subset within IL-21R+ | group1 | group2 | n1 | n2 | statistic | df | p |
|---|---|---|---|---|---|---|---|---|
| Mock | Naive | IL21 | NT | 10 | 10 | -3.96 | 9 | 0.00331 |
| KSHV | Plasma Cell | IL21 | NT | 10 | 10 | 2.39 | 9 | 0.0405 |
| KSHV | Plasmablast | IL21 | NT | 10 | 10 | 2.10 | 9 | 0.0647 |
| Mock | Plasmablast | IL21 | NT | 10 | 10 | 1.76 | 9 | 0.112 |
| KSHV | Double Neg | IL21 | NT | 10 | 10 | -1.68 | 9 | 0.127 |
| KSHV | MZ-like | IL21 | NT | 10 | 10 | -1.53 | 9 | 0.161 |
| Mock | MZ-like | IL21 | NT | 10 | 10 | -1.48 | 9 | 0.173 |
| Mock | Plasma Cell | IL21 | NT | 10 | 10 | 1.41 | 9 | 0.191 |
| KSHV | Naive | IL21 | NT | 10 | 10 | -1.06 | 9 | 0.316 |
| KSHV | Germinal Center | IL21 | NT | 10 | 10 | -0.75 | 9 | 0.47 |
| KSHV | Memory | IL21 | NT | 10 | 10 | 0.71 | 9 | 0.498 |
| Mock | Memory | IL21 | NT | 10 | 10 | -0.66 | 9 | 0.526 |
| Mock | Double Neg | IL21 | NT | 10 | 10 | 0.52 | 9 | 0.614 |
| Mock | Germinal Center | IL21 | NT | 10 | 10 | -0.39 | 9 | 0.706 |
| KSHV | Transitional | IL21 | NT | 10 | 10 | -0.37 | 9 | 0.719 |
| Mock | Transitional | IL21 | NT | 10 | 10 | 0.33 | 9 | 0.746 |

**Supplemental Table 4D**: Two-way repeated measures ANOVA analysis for the effect of IL-21 treatment (TX) and Baseline vs. Mock vs. KSHV infection (Cond) on the MFI of IL-21R within IL-21R+ B cell subsets at 3 dpi. Sorted by p value.

| **MFI of IL-21R at 3dpi** | Effect | DFn | DFd | F | p | p<.05 | ges |
|---|---|---|---|---|---|---|---|
| CD20+ B cell | Cond | 1 | 9 | 4.666 | 0.059 | | 0.024 |
| Transitional | Cond:TX | 1 | 9 | 4.587 | 0.061 | | 0.003 |
| CD20- Plasma Cell | Cond | 1 | 9 | 3.634 | 0.089 | | 0.054 |
| Plasmablast | TX | 1 | 9 | 3.469 | 0.095 | | 0.032 |
| Double Neg | Cond:TX | 1 | 9 | 3.337 | 0.101 | | 0.071 |
| CD20+ B cell | TX | 1 | 9 | 3.283 | 0.103 | | 0.035 |
| CD20- Plasma Cell | Cond:TX | 1 | 9 | 3.236 | 0.106 | | 0.113 |
| Naive | TX | 1 | 9 | 3.229 | 0.106 | | 0.034 |
| Transitional | TX | 1 | 9 | 2.826 | 0.127 | | 0.011 |
| MZ-like | TX | 1 | 9 | 2.722 | 0.133 | | 0.069 |
| Centroblast | Cond | 1 | 9 | 2.533 | 0.146 | | 0.038 |
| CD20- Plasma Cell | TX | 1 | 9 | 2.515 | 0.147 | | 0.043 |
| Plasma Cell | Cond:TX | 1 | 9 | 2.45 | 0.152 | | 0.064 |
| Transitional | Cond | 1 | 9 | 2.267 | 0.166 | | 0.013 |
| CD20+ Plasma Cells | Cond:TX | 1 | 9 | 2.194 | 0.173 | | 0.03 |
| Plasma Cell | Cond | 1 | 9 | 2.162 | 0.176 | | 0.058 |

| | | | | | | | |
|---|---|---|---|---|---|---|---|
| Centrocyte | TX | 1 | 9 | 1.929 | 0.198 | | 0.044 |
| CD20+ Plasma Cells | Cond | 1 | 9 | 1.905 | 0.201 | | 0.075 |
| Germinal Center | TX | 1 | 9 | 1.868 | 0.205 | | 0.055 |
| Plasma Cell | TX | 1 | 9 | 1.756 | 0.218 | | 0.05 |
| MZ-like | Cond | 1 | 9 | 1.378 | 0.271 | | 0.039 |
| CD20+ Plasma Cells | TX | 1 | 9 | 1.346 | 0.276 | | 0.02 |
| Germinal Center | Cond:TX | 1 | 9 | 1.258 | 0.291 | | 0.023 |
| MZ-like | Cond:TX | 1 | 9 | 1.229 | 0.296 | | 0.035 |
| Germinal Center | Cond | 1 | 9 | 1.054 | 0.331 | | 0.02 |
| CD20- B cells | Cond:TX | 1 | 9 | 0.908 | 0.366 | | 0.031 |
| Naive | Cond:TX | 1 | 9 | 0.865 | 0.377 | | 0.032 |
| Naive | Cond | 1 | 9 | 0.738 | 0.413 | | 0.015 |
| CD20- B cells | Cond | 1 | 9 | 0.73 | 0.415 | | 0.014 |
| Memory | TX | 1 | 9 | 0.553 | 0.476 | | 0.003 |
| CD20- B cells | TX | 1 | 9 | 0.463 | 0.513 | | 0.01 |
| Plasmablast | Cond | 1 | 9 | 0.339 | 0.575 | | 0.004 |
| CD20+ B cell | Cond:TX | 1 | 9 | 0.282 | 0.608 | | 0.002 |
| Plasmablast | Cond:TX | 1 | 9 | 0.213 | 0.655 | | 0.003 |
| Centrocyte | Cond:TX | 1 | 9 | 0.19 | 0.673 | | 0.006 |
| Double Neg | Cond | 1 | 9 | 0.126 | 0.731 | | 0.004 |
| Centrocyte | Cond | 1 | 9 | 0.111 | 0.746 | | 0.004 |
| Memory | Cond:TX | 1 | 9 | 0.088 | 0.774 | | 0.001 |
| Centroblast | TX | 1 | 9 | 0.083 | 0.779 | | 0.002 |
| Memory | Cond | 1 | 9 | 0.026 | 0.875 | | 0.000513 |
| Double Neg | TX | 1 | 9 | 0.001 | 0.971 | | 3.37e-05 |
| Centroblast | Cond:TX | 1 | 9 | 0.001 | 0.973 | | 4.45e-05 |

**Supplemental Table 5A**: Pairwise correlations using Pearson's method between T cell frequencies within IL-21+ T cells and total GFP+ B lymphocytes at 3 dpi. Sorted by p value.

| **T cell subset within IL-21+** | **Cor to total GFP+** | statistic | p | p<.05 | conf.low | conf.high |
|---|---|---|---|---|---|---|
| CD8_CM | 0.57 | 2.42 | 0.03 | * | 0.06 | 0.85 |
| CD4_TM | -0.51 | -2.05 | 0.06 | | -0.82 | 0.03 |
| CD4_EM | -0.45 | -1.72 | 0.11 | | -0.79 | 0.11 |
| CD4_RO | -0.45 | -1.72 | 0.11 | | -0.79 | 0.11 |
| CD4 | -0.41 | -1.57 | 0.14 | | -0.77 | 0.15 |
| CD8_SCM | -0.41 | -1.54 | 0.15 | | -0.77 | 0.16 |
| CD8_RO | 0.4 | 1.52 | 0.16 | | -0.16 | 0.77 |
| CD8_RA | -0.39 | -1.46 | 0.17 | | -0.76 | 0.18 |
| CD4_CM | -0.36 | -1.35 | 0.2 | | -0.75 | 0.21 |
| CD8_TEMRA | -0.35 | -1.31 | 0.21 | | -0.75 | 0.22 |
| CD8_Treg | -0.35 | -1.31 | 0.21 | | -0.74 | 0.22 |
| CD4_Naive | 0.31 | 1.13 | 0.28 | | -0.26 | 0.72 |
| CD4_Th17 | 0.29 | 1.05 | 0.31 | | -0.28 | 0.71 |
| CD8_TM | -0.27 | -0.98 | 0.35 | | -0.7 | 0.3 |

| | | | | | | |
|---|---|---|---|---|---|---|
| CD4_BCL6- TfH | -0.26 | -0.92 | 0.37 | | -0.69 | 0.32 |
| CD8_EM | -0.26 | -0.92 | 0.38 | | -0.69 | 0.32 |
| CD8_Tc17 | 0.25 | 0.88 | 0.4 | | -0.33 | 0.69 |
| CD4_Tfh | -0.23 | -0.83 | 0.42 | | -0.68 | 0.34 |
| CD4_Treg | -0.22 | -0.8 | 0.44 | | -0.67 | 0.35 |
| CD4_BCL6+ TfH | -0.22 | -0.78 | 0.45 | | -0.67 | 0.35 |
| CD4_TE | -0.2 | -0.72 | 0.48 | | -0.66 | 0.37 |
| CD4_SCM | 0.18 | 0.65 | 0.53 | | -0.38 | 0.65 |
| CD4_RA | 0.17 | 0.59 | 0.56 | | -0.4 | 0.64 |
| CD4_TEMRA | 0.16 | 0.55 | 0.59 | | -0.41 | 0.64 |
| CD8 | -0.15 | -0.52 | 0.61 | | -0.63 | 0.41 |
| CD8_TE | 0.12 | 0.41 | 0.69 | | -0.44 | 0.61 |
| IL21 | 0.1 | 0.33 | 0.74 | | -0.46 | 0.6 |
| CD8_Naive | 0.01 | 0.03 | 0.98 | | -0.52 | 0.54 |

**Supplemental Table 5B**: Pairwise correlations using Pearson's method between the iMFI of IL-21 for T cell subsets and total GFP+ B lymphocytes at 3 dpi. Sorted by p value.

| **iMFI of IL-21 for T cell subset** | **Cor to total GFP** | statistic | p | p<.05 | conf.low | conf.high |
|---|---|---|---|---|---|---|
| CD4_TE | -0.25 | -1.31 | 0.2 | | -0.57 | 0.14 |
| CD8_TEMRA | -0.23 | -1.21 | 0.24 | | -0.56 | 0.16 |
| CD8_RA | -0.19 | -0.99 | 0.33 | | -0.53 | 0.2 |
| CD8_TM | -0.17 | -0.89 | 0.38 | | -0.51 | 0.21 |
| CD4_TM | -0.16 | -0.83 | 0.42 | | -0.5 | 0.23 |
| CD4_TEMRA | 0.16 | 0.83 | 0.42 | | -0.23 | 0.5 |
| CD4 | -0.14 | -0.7 | 0.49 | | -0.48 | 0.25 |
| CD4_RO | -0.12 | -0.64 | 0.53 | | -0.48 | 0.26 |
| CD4_Tfh | -0.11 | -0.58 | 0.57 | | -0.47 | 0.27 |
| CD8_CM | 0.11 | 0.57 | 0.58 | | -0.27 | 0.46 |
| CD8_Tc17 | -0.11 | -0.56 | 0.58 | | -0.46 | 0.27 |
| CD8_RO | -0.1 | -0.5 | 0.62 | | -0.45 | 0.29 |
| CD4_BCL6+ TfH | -0.1 | -0.48 | 0.63 | | -0.45 | 0.29 |
| CD8_TE | 0.08 | 0.41 | 0.68 | | -0.3 | 0.44 |
| CD4_Treg | -0.08 | -0.41 | 0.69 | | -0.44 | 0.3 |
| CD4_CM | -0.08 | -0.4 | 0.7 | | -0.44 | 0.3 |
| CD4_EM | -0.08 | -0.39 | 0.7 | | -0.44 | 0.31 |
| CD8_SCM | -0.06 | -0.33 | 0.74 | | -0.43 | 0.32 |
| CD8_EM | 0.06 | 0.31 | 0.76 | | -0.32 | 0.42 |
| CD8_Na | -0.05 | -0.27 | 0.79 | | -0.42 | 0.33 |
| CD8_Treg | 0.05 | 0.24 | 0.81 | | -0.33 | 0.41 |
| CD4_Na | 0.04 | 0.22 | 0.83 | | -0.34 | 0.41 |
| CD4_SCM | -0.04 | -0.18 | 0.86 | | -0.4 | 0.34 |
| CD4_RA | -0.03 | -0.17 | 0.86 | | -0.4 | 0.34 |
| CD8 | -0.03 | -0.17 | 0.87 | | -0.4 | 0.34 |
| CD4_BCL6- TfH | -0.02 | -0.09 | 0.93 | | -0.39 | 0.36 |

| | | | | | | |
|---|---|---|---|---|---|---|
| CD4_Th17 | 0 | 0 | 1 | | -0.37 | 0.37 |

**Supplemental Table 5C**: Pairwise correlations using Pearson's method between the baseline (Day 0) frequencies T cell subsets and total GFP+ B lymphocytes at 3 dpi. Sorted by p value.

| **Baseline frequency of T cell subset** | **Cor to total GFP+** | statistic | p | p<.05 | conf.low | conf.high |
|---|---|---|---|---|---|---|
| CD4_TE | 0.58 | 2.46 | 0.03 | * | 0.07 | 0.85 |
| CD4_RO | -0.57 | -2.4 | 0.03 | * | -0.84 | -0.06 |
| CD8_SCM | -0.47 | -1.82 | 0.09 | | -0.8 | 0.09 |
| CD8_RO | 0.47 | 1.86 | 0.09 | | -0.08 | 0.8 |
| CD4_Naive | 0.46 | 1.81 | 0.1 | | -0.09 | 0.8 |
| CD8_TE | -0.46 | -1.8 | 0.1 | | -0.8 | 0.09 |
| CD8_EM | 0.44 | 1.72 | 0.11 | | -0.11 | 0.79 |
| CD8_CM | 0.43 | 1.66 | 0.12 | | -0.13 | 0.78 |
| CD4_TM | -0.41 | -1.54 | 0.15 | | -0.77 | 0.16 |
| CD8_TM | 0.39 | 1.45 | 0.17 | | -0.18 | 0.76 |
| CD4_Tfh | -0.38 | -1.44 | 0.18 | | -0.76 | 0.18 |
| CD8 | -0.37 | -1.4 | 0.19 | | -0.75 | 0.19 |
| CD8_Tc17 | 0.36 | 1.32 | 0.21 | | -0.21 | 0.75 |
| CD4_RA | 0.31 | 1.14 | 0.28 | | -0.26 | 0.72 |
| CD4 | 0.25 | 0.9 | 0.38 | | -0.32 | 0.69 |
| CD4_TEMRA | 0.22 | 0.78 | 0.45 | | -0.35 | 0.67 |
| CD4_Th17 | 0.19 | 0.65 | 0.52 | | -0.38 | 0.65 |
| CD4_Treg | -0.15 | -0.53 | 0.61 | | -0.63 | 0.41 |
| CD8_TEMRA | -0.15 | -0.52 | 0.62 | | -0.63 | 0.42 |
| CD4_CM | -0.13 | -0.46 | 0.65 | | -0.62 | 0.43 |
| CD4_BCL6-1 | -0.13 | -0.47 | 0.65 | | -0.62 | 0.43 |
| CD4_BCL6+ | 0.12 | 0.4 | 0.7 | | -0.44 | 0.61 |
| CD8_RA | -0.1 | -0.35 | 0.73 | | -0.6 | 0.45 |
| CD8_Treg | 0.08 | 0.27 | 0.8 | | -0.47 | 0.58 |
| CD4_SCM | -0.06 | -0.22 | 0.83 | | -0.58 | 0.48 |
| CD4_EM | -0.05 | -0.19 | 0.86 | | -0.57 | 0.49 |
| CD8_Naive | -0.04 | -0.14 | 0.89 | | -0.56 | 0.5 |
